# PD-L1 lncRNA splice promotes lung adenocarcinoma progression via enhancing c-Myc activity

**DOI:** 10.1101/2020.09.29.282541

**Authors:** Shuang Qu, Zichen Jiao, Geng Lu, Bing Yao, Ting Wang, Weiwei Rong, Jiahan Xu, Ting Fan, Xinlei Sun, Rong Yang, Jun Wang, Yongzhong Yao, Guifang Xu, Xin Yan, Tao Wang, Hongwei Liang, Ke Zen

## Abstract

Although blockade of programmed death-ligand 1 (PD-L1) to enhance T cell immune responses shows great promise in tumor immunotherapy, the efficacy of such immune-checkpoint inhibition strategy is limited for patients with solid tumors. The mechanism underlying the limited efficacy of PD-L1 inhibitors remains unclear. Here, we show that human lung adenocarcinoma, regardless of PD-L1 protein positive or negative, all produce a long non-coding RNA isoform of PD-L1 (PD-L1-lnc) via alternative splicing, which promotes lung adenocarcinoma proliferation and metastasis. PD-L1-lnc in various lung adenocarcinoma cells is significantly upregulated by IFNγ in a manner similar to PD-L1 mRNA. Both *in vitro* and *in vivo* studies demonstrate that PD-L1-lnc increases proliferation and invasion but decreases apoptosis of lung adenocarcinoma cells. Mechanistically, PD-L1-lnc directly binds to c-Myc and enhances c-Myc transcriptional activity downstream in lung adenocarcinoma cells. Our results provide targeting PD-L1-lnc−c-Myc axis as a novel strategy for lung cancer therapy.

## INTRODUCTION

Cancer is one of the major threats to human health worldwide and is responsible for millions of deaths annually [1]. Cancer cells have developed sophistic mechanisms to evade immune surveillance and attack by immune cells. One such mechanism is to express various immune checkpoint molecules, which suppress anti-tumor immunity of immune cells. Among these intrinsic negative checkpoints, programmed death protein 1 (PD-1) and its ligand 1 (PD-L1) are prominent pair and the blockade of PD-1/PD-L1 has led to successful immunotherapies. PD-L1, also known as B7-H1 and CD274, is a transmembrane protein commonly expressed on the surface of tumor cells. PD-L1 specifically binds to its receptor, PD-1, which is expressed on the surface of T cells, B cells and myeloid cells [2]. The binding of PD-L1 to PD-1 is able to activate the downstream signaling of PD-1 receptor in T cells, resulting in inhibition of the proliferation, cytokine generation/release and cytotoxicity of T cells. The existence of a break on T cell function may prevent autoimmunity, but many tumor cells exploit this mechanism to protect themselves from immune attack, resulting in tumor immune evasion [3]. The commercial PD-L1 antibodies have shown tremendous success in treating melanoma and blood cancers like leukemia and lymphoma [4, 5]. However, the efficacy of PD-L1 inhibition is limited for many cancer patients with solid tumors [6–8]. The molecular basis underlying low efficacy of PD-L1 blockade in solid tumor is unclear though several contributing factors have been suggested. First, IFNγ fails to induce the expression of PD-L1 protein in certain cancers [9, 10]. It has been shown that PD-L1 can express in tumor cells as different forms, including membrane PD-L1 (mPD-L1), cellular PD-L1 (cPD-L1) and soluble PD-L1 (sPD-L1)[11, 12]. Given that only mPD-L1 can bind to T cell surface PD-1 and suppress T cell function, low level of mPD-L1 may result in poor efficacy of anti-PD-L1 antibody. These PD-L1 negative tumors had been characterized as ‘cold cancer’ as opposed to PD-L1 positive tumors, and indeed cold tumors showed low efficacy of PD-L1 blockade treatment [13]. An additional factor contributing limited efficacy of PD-L1 blockade is a lack of infiltration of anti-tumor T cells into tumor microenvironment (TME) due to poor initial antigen presentation [14, 15]. As PD-L1 executes its function via targeting T cells, low level of T cell infiltration would be associated with limited efficacy of PD-L1 blockade.

Long non-coding RNAs (lncRNAs) have recently gained attention in the field of cancer research [16–18]. As RNA transcripts of 200 or more nucleotides that are not translated to proteins, lncRNAs can be transcribed from their own promotors, from the promotors of other coding or non-coding sequences of DNA or from the enhancer sequences. Some lncRNAs are derived through alternative splicing of transcribed RNA, a process which occurs in over 90% of human multi-exon protein-coding genes [19]. Several classes of lncRNAs have been discovered on the basis of diverse parameters such as transcript length, association with annotated protein-coding genes and mRNA resemblance among others [20]. Recent discoveries suggest that lncRNAs are relevant to cancer progression whereby lncRNAs interact with both oncogenic and tumor suppressive pathways [16]. In line with this, the expression of lncRNAs has been widely reported to be dysregulated in various human cancers [17, 18]. Despite these findings, the biogenesis and regulatory role of lncRNAs in cancer development, however, remains incompletely understood.

In the present study, we report that, in addition to PD-L1 mRNA, human lung adenocarcinoma PD-L1 gene can generate a long non-coding RNA (PD-L1-lnc) through alternative splicing. Moreover, in a similar manner to PD-L1 mRNA, PD-L1-lnc in lung adenocarcinoma is markedly upregulated by IFNγ. Once generated, PD-L1-lnc promotes lung adenocarcinoma cell proliferation and invasion but decreases cell apoptosis via directly binding to c-Myc and activating c-Myc transcriptional activity. Taken together, this study identifies PD-L1-lnc−c-Myc axis as a novel mechanism underlying human lung adenocarcinoma progression.

## MATERIALS AND METHODS

### Clinical samples

The 275 pairs of lung adenocarcinoma tissues and adjacent non-cancerous tissues were collected from patients who were diagnosed with lung adenocarcinoma at the Nanjing Drum Tower Hospital (Nanjing, China) and who had not yet received treatment. The clinic pathological features are described in Table S1. Tissues were collected after surgical resection and stored in liquid nitrogen before use. The study was authorized by the Ethics Committee of the Nanjing Drum Tower Hospital. All experiments were performed in accordance with relevant guidelines and regulations.

### Cell lines, culture conditions and IFNγ treatment

Human lung cancer cell lines A549, PC9, H1975, H1299 and H1650 were purchased from American Type Culture Collection (ATCC) (Manassas, VA, USA). Cells were cultured in Dulbecco’s Modified Eagle’s Medium (DMEM, Gibco, Carlsbad, CA, USA) supplemented with 10% (v/v) FBS (Gibco, Carlsbad, CA, USA), 1% (v/v) penicillin/streptomycin (Gibco, Carlsbad, CA, USA). For IFNγ treatment, cells were seeded into six-well plates on day 1, targeting 70%–80% of confluence on the day of surface staining. On day 2, cells were exposed to 100ng/mL IFNγ (PeproTech, USA) for 24 h.

### Immunohistochemical staining

Tissues were fixed in 4% paraformaldehyde, embedded in paraffin, and cut into 4μm sections. Immunohistochemical staining of PD-L1 was performed using the anti-PD-L1 antibody (SP142, Zhongshanjinqiao, China) according to the manufacturer’s protocol.

### RNA extraction and RT-PCR/qRT-PCR assay

Total RNAs were isolated by TRIzol™ Reagent (Invitrogen, US) according to the manufacturer’s instruction. The quality and quantity of RNA was assayed by Nanodrop 2000 spectrophotometer (Thermo Fisher Scientific, USA). The nuclear and cytoplasmic fractions were purified by PARIS Kit (Ambion, Life Technologies, USA). RNA was reverse transcribed by HiScript II Q RT SuperMixfor qPCR (+gDNA wiper) (Vazyme, Nanjing, China). The 1.1×T3 Super PCR Mix (TsingKe, Nanjing, China) was used for PCR. The 10% vertical polyacrylamide electrophoresis was performed to observe the cDNA PCR products. AceQ qPCR SYBR Green Master Mix (Vazyme, Nanjing, China) was used for qRT-PCR, and GAPDH was used to normalize the level of PD-L1-lnc and PD-L1 mRNA. Primers were listed in table S2. The purified fragments were cloned into the pCR® II TA vector using the TA cloning kit (Thermo Fisher Scientific, United States) and sequenced at the TsingKe Biotechnology (Nanjing, China) to validate the cDNA PCR products.

### RNA BaseScope

BaseScope™ Probe for PD-L1-lnc and PD-L1 mRNA were designed and synthesized by ACD (Cat. No.700001, 700001-C2, Advanced Cell Diagnostics, CA, USA). Tissues and cells were fixed by 10% neutral buffered formalin on slides for detection of PD-L1-lnc and PD-L1 mRNA. The signals of the PD-L1-lnc and PD-L1 mRNA probes were detected by BaseScope™ Detection Reagent Kit (ACD, USA) according to the manufacturer’s instructions. The images were acquired on Leica SP5 Scanning Laser Confocal Microscope (Leica Microsystems, Wetzlar, Germany).

### Flow cytometry

Flow cytometry was performed using a CytoFLEX S (Beckman Coulter Life Sciences, Mississauga, ON). Human lung adenocarcinoma cell lines A549, PC9, H1299, H1650 and H1975 were stained with 0.2µg of PE-conjugated anti-PD-L1 (Biolegend, USA) antibody.

### Western blotting

The protein extraction reagent (Thermo Scientific) with a cocktail of proteinase inhibitors (Roche Applied Science, Switzerland) was used to isolate the total protein from cells or tissue samples. Equal amount of total protein was separated by 10% SDS–PAGE and transferred onto a PVDF membrane. Then, the membranes were blocked with 5% skimmed milk and incubated the membranes with primary antibodies at 4°C overnight and then incubated with secondary antibodies at room temperature for 2 h. The bands were examined by Immobilob™ Western Chemiluminescent HRP Substrate (Millipore, Billerica, MA, USA). The primary antibody and secondary antibodies detailed information list below: PD-L1 [E1L3N®] XP ® Rabbit mAb (#13684, Cell Signaling Technology, Beverly, MA, USA), GAPDH (6C5) Mouse mAb (sc-32233, Santa Cruz, USA), Histone-H3 Rabbit Polyclonal antibody (17168-1-AP, Proteintech, Wuhan, China), c-MYC Rabbit Polyclonal antibody (10828-1-AP, Proteintech, Wuhan, China), MAX antibody [EPR19352] (ab199489, abcam, US), Rabbit (DA1E) mAb IgG XP® Isotype Control (#3900, Cell Signaling Technology, Beverly, MA, USA) and the secondary antibodies (goat anti-rabbit IgG-HRP, sc-2030, Santa Cruz; goat anti-mouse IgG-HRP, sc-2005, Santa Cruz; Mouse Anti-rabbit IgG-HRP [L27A9] mAb, #5127, Cell Signaling Technology).

### Cell proliferation assay

To measure the proliferation rate of human lung cancer cells, EdU assays were performed. Briefly, A549 and PC9 cells were seeded in 6-well plates and transfected with vectors. At 24 h after transfection, cells were harvested and reseeded 48-well plates for EdU assays. The EdU assay kit (RiBoBio, China) was used to determine the proliferation rate of cells according to the manufacturer’s instructions.

### Cell invasion assay

Firstly, the upper chamber of transwell (Millipore, USA) was coated with diluted Matrigel (200 mg/mL, BD Biosciences, MA) at the density of 50 µL/well before use. A549 and PC9 cells were transfected with vectors. After 24 h, cells were resuspended in FBS-free DMEM medium and reseeded on the upper surface of 24-well plates. Cells were allowed to migrate across the 8-μm membrane toward medium with 20% FBS for 24 h. Then, the cells were fixed with 4% paraformaldehyde and dyed with 0.5% crystal violet. Non-migrating cells were removed using a cotton swab. The migrant cells were blindly counted under a light microscope (Leica Microsystems, Wetzlar, Germany).

### Cell apoptosis assay

The apoptosis of A549 and PC9 cells was assayed using the Annexin V-Alexa Fluor 647/PI (YEASEN, China) based on the procedures provided by the manufacturer. The transfected A549 and PC9 cells were cultured in serum-free DMEM for 24 h. The collected cells were washed with cold PBS and re-suspended in 1 × binding buffer, followed by staining with Alexa Fluor 647-Annexin V and propidium iodide (PI) in the dark for 15 min. The apoptotic cells were calculated using flow cytometer (Beckman Coulter Life Sciences, Mississauga, ON).

### Xenograft assays in nude mice

All animal care and handling procedures were performed in accordance with the National Institutes of Health’s Guide for the Care and Use of Laboratory Animals. Male athymic BALB/c nude mice (6 weeks old) were purchased from the Model Animal Research Center of Nanjing University (Nanjing, China) and were randomly divided into 3 groups (5 mice per group). A549 cells were firstly transfected with control vector, vector expressing PD-L1-lnc (PD-L1-lnc) or PD-L1-lnc shRNA (PD-L1-lnc shRNA), respectively. Cells were then selected using 500µg/mL G418 (ThermoFisher) or 1µg/mL Puromycin (ThermoFisher) for 2 weeks. Three stable lung cancer cell lines were then subcutaneously injected into mice (10^6^ cells/0.1 ml PBS per mouse). The needle was inserted into the armpit of the left foreleg at a 45 degree angle and a 5 mm depth, midway down. Then, the longest diameter (a) and the shortest diameter (b) of the tumor were measured using digital calipers every three days, and the tumor volume (V) was calculated according to formula: V=a×b^2^/2. Mice were sacrificed and photographed 15 days post-injection. The xenograft tumors were removed and analyzed. Tissues were subjected to extraction of total RNA, as well as hematoxylin and eosin (H&E) staining and Ki67 immunohistochemical staining.

### Vector construction and cell transfection

To overexpress PD-L1-lnc, the full-length cDNA of PD-L1-lnc was synthesized and cloned into pcDNA3.1-P2A-eGFP vector (GenScript, China). To suppress PD-L1-lnc, the PD-L1-lnc shRNA vectors were synthesized and then cloned into pLKO.1 vector (GenScript, China). The siRNA target sequences were listed in table S3. Cells were transfected using Lipofectamine 3000 (Invitrogen, USA) according to the manufacturer’s instruction.

### Microarray analysis

Total RNAs were isolated from the A549 cells transfected with PD-L1-lnc overexpressed vector or control vector by TRIzol reagent and purified by RNeasy Mini Kit (Qiagen, USA). RNA samples were performed to Microarray analysis by Agilent SurePrint G3 human gene expression Microarray 8X60K (Agilent Technologies). After hybridization, slides were scanned on the Agilent Microarray Scanner. Data were extracted using Feature Extraction software 10.7 (Agilent Technologies). Raw data were normalized by Quantile algorithm, limma package the R program. Significant differential expressed transcripts were defined as fold change ≥2 or ≤−2 and P-value ≤0.05.

### Streptavidin pull down of PD-L1-lnc and c-Myc protein

For each pull down sample 100μl of streptavidin magnetic beads (S1420S, New England Biolabs, US) were washed with wash/binding buffer (0.5 M NaCl, 20 mM Tris-HCl, pH 7.5, 1 mM EDTA) twice, then incubation with probes for 1 hour at 4℃. For biotin-coupled RNA capture, the 5’-end biotinylated PD-L1-lnc probe or control RNA were used. Probes used as follows: PD-L1-lnc probe: 5’-3’ CATCCATCATTCTCCCAAGTGAGTCCT, GFP probe: 5’-3’ TGAAGTTCACCTTGATGCCGTTCTTCT GCTTGTCGGCCATGATAT AGACGTTGTGGCTGT. The 0.5 ml cell lysis buffer (Invitrogen, USA) with complete protease inhibitor cocktail (Roche Applied Science, IN) and Recombinant RNase Inhibitor (Takara, Japan) were added into the cell pellets, and lysed by sonication. The cell lysates incubation with RNA-coupled beads followed by centrifugation at 13,000×rpm for 20 min. After rotating for 4 h at 4°C, the beads were washed with 1 ml lysis buffer (containing 300 mM NaCl) twice, 1 ml low-salt lysis buffer (containing 150 mM NaCl). Half of beads were resuspended in TRIzol™ Reagent for detecting PD-L1-lnc level, while half of beads were resuspended in RIPA lysis buffer for detecting c-Myc protein level.

### RNA immunoprecipitation (RIP)

A549 Cells transfected with PD-L1-lnc overexpressed vector or control vector were lysed with lysis buffer (20 mM Tris-HCl, 150 mM NaCl, 0.5% Nonidet P-40, 2 mM EDTA, 0.5 mM DTT, 1 mM NaF, 1 mM PMSF and 1% Protease Inhibitor Cocktail from Sigma, pH 7.4) for 30 min on ice. After cleared by centrifugation (16,000g) for 10 min at 4 °C, the lysates were subjected to immunoprecipitation with anti-cMyc antibody or IgG followed by protein A/G-Agarose beads. After the elution, the proteins were isolated by RIPA lysis buffer for western blot assays and the RNA were isolated with TRIzol™ Reagent for detecting PD-L1-lnc level.

### *In vitro* mRNA stability assay

To monitor the effect of PD-L1-lnc on the stability of PD-L1 mRNA, A549 cells transfected with control vector and PD-L1-lnc vector were treated by 5μg/ml actinomycin D to block the transcription of new PD-L1 mRNA, and then assessed PD-L1 mRNA level at 0, 3, 6, 9, 12 or 24 h post-treatment by qRT-PCR.

### Statistical analysis

All experiments were performed in triplicate or as indicated in the experiments. Data were presented as the mean ± SD. When only two groups were compared, Student’s *t*-test was used. Comparisons involving multiple dependent measures were Tukey-Kramer corrected. The reported *P* value was 2-sided. The differences with **P<0.05 were considered significantly different.

## RESULTS

### Generation of PD-L1 lncRNA via alternative splicing in PD-L1 protein positive or negative human lung adenocarcinoma tissues, as well as various lung adenocarcinoma cells

By immunohistochemical staining, we screened more than 275 human lung adenocarcinoma samples and their paired distal non-cancerous tissues samples from patients registered in Nanjing Drum Tower Hospital, Nanjing University School of Medicine (Nanjing, Jiangsu, China) from 2017-2018 (Table S1) for PD-L1 protein expression. As shown in fig. S1A and Table S1, the majority of lung cancer samples were PD-L1 negative, with little or no PD-L1 protein expression, which is in line with the previous report about the PD-L1 protein expression in Chinese lung cancer patients [21]. Although the percentage of PD-L1-positive lung cancers at advanced stage was higher than that of lung cancers at early stage [22], considerable numbers of lung adenocarcinoma samples at advanced stages expressed little or no PD-L1 protein (fig. S1A and Table S1).

To explore the mechanism underlying the variety of PD-L1 protein expression in lung adenocarcinoma at various stages, we detected the PD-L1 mRNA level in PD-L1 protein-positive or PD-L1 protein-negative lung cancer tissues. To our surprise, analysis of PCR end-product identified two bands at 198bp and 92bp, respectively (Fig. 1A, left). Sanger sequencing showed that 198bp band matched to the sequence of PD-L1 mRNA (Fig. 1A, right upper), while 92bp band was a noncoding isoform (NR_052005.1) which lacks an alternate internal segment (named as PD-L1-lnc) (Fig. 1A, right lower). To confirm these results, another pair of probes for detecting the two-missing parts was designed (Fig. 1B, upper). The results of agarose gel showed two bands at 878bp and 705bp, respectively (Fig. 1B, lower left). The 878bp band matched the sequence of PD-L1 mRNA (Fig. 1B, upper right; fig. S2A), while the 705bp band matched to the NR_052005.1 missing 106nt in exon 4 and 67nt between exon 5 and 6 (Fig. 1B, lower right; fig. S2B). Furthermore, RNA BaseScope analysis [23] was performed to examine the co-expression of PD-L1 mRNA and PD-L1-lnc in human lung cancer tissues. As shown in Fig. 1C, a clear co-existence of PD-L1 mRNA (blue dots) and PD-L1-lnc (red dots) in lung cancer tissues was observed. To accurately quantify these RNA fragments, the specific probes for PD-L1 mRNA and PD-L1-lnc were designed (Fig. 1D, upper). The PCR products of these probes (for PD-L1 mRNA and PD-L1-lnc) were confirmed by agarose gel (Fig. 1D, lower left) and Sanger sequencing (fig. S3A-B), respectively. The qRT-PCR assay with specific probes detected similar amount of PD-L1 mRNA (Fig. 1D, lower middle) and PD-L1-lnc (Fig. 1D, lower right) in PD-L1 protein-positive and PD-L1 protein-negative tumor samples.

**Figure 1.**
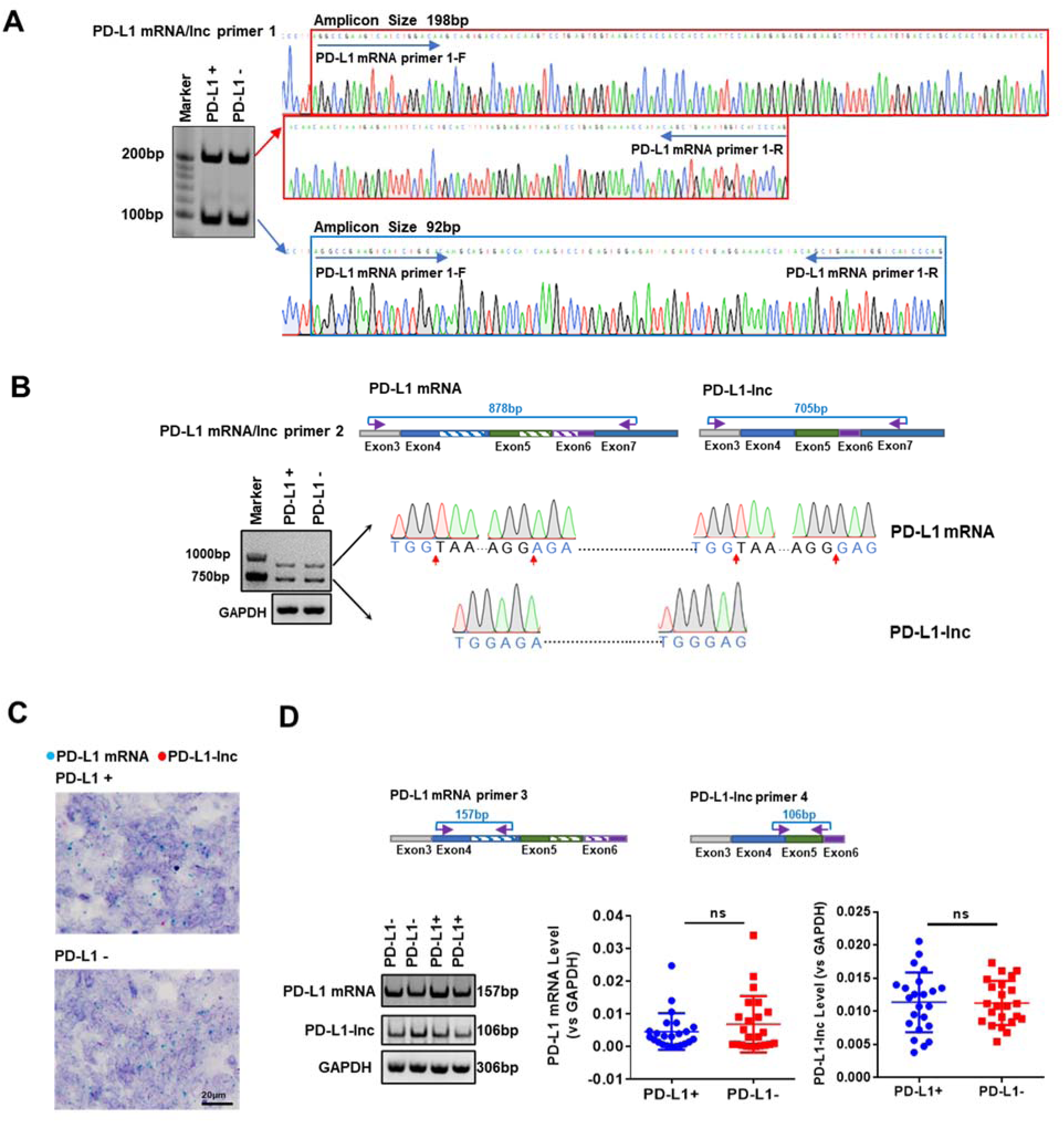
Generation of PD-L1-lnc in PD-L1 protein positive or negative lung adenocarcinoma via alternative splicing. **(A)** Agarose gel analysis and Sanger sequencing of the PCR end-product in PD-L1 protein positive or negative tumor tissues. **(B)** Upper, the schematic of primers for amplification of PD-L1 mRNA and PD-L1-lnc; Lower, the agarose gel and Sanger sequencing of the 878bp and 705bp band obtained by RT-PCR. **(C)** RNA BaseScope for PD-L1 mRNA (blue) and PD-L1-lnc (red) in PD-L1 protein positive and negative tumor tissues. **(D)** Upper, the schematic of primer for amplification of PD-L1 mRNA and PD-L1-lnc; Lower, the expression level of PD-L1 mRNA and PD-L1-lnc in PD-L1 protein-positive or -negative tumor tissues by qRT-PCR with specific probes.

Expression of PD-L1-lnc and PD-L1 mRNA was next detected in various human lung cancer cell lines, including A549, PC9, H1975, H1650 and H1299. Western blotting and flow cytometry analysis confirmed various levels of PD-L1 on the lung adenocarcinoma cell lines (Fig. 2A-B). Similar to that in human lung adenocarcinoma tissues, both PD-L1-lnc and PD-L1 mRNA were detected in various lung cancer cells in various human lung cancer cells by agarose gel through PD-L1 mRNA/lnc primer 2 (Fig. 2C), RNA BaseScope (Fig. 2D) and RT-PCR by PD-L1 mRNA specific primer 3 or PD-L1-lnc specific primer 4 (Fig. 2E).

**Figure 2.**
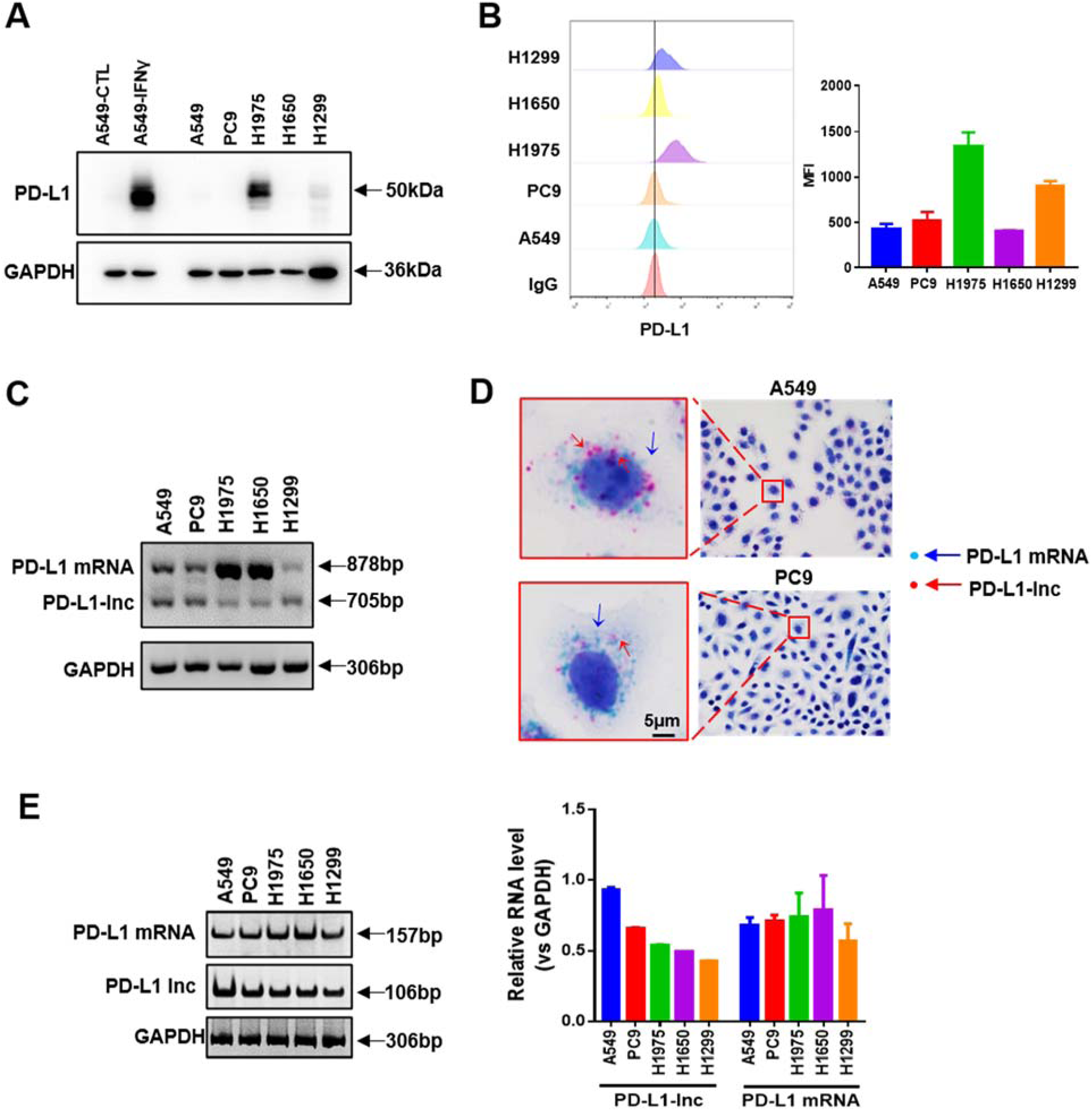
The expression of PD-L1-lnc in various lung adenocarcinoma cancer cell lines. **(A-B)** Western blotting (A) and flow cytometry (B) analysis of PD-L1 in various lung adenocarcinoma cancer cell lines. **(C)** Agarose gel analysis of PCR end-product of PD-L1 transcript showing two bands at 878bp and 705bp. **(D)** RNA BaseScope for PD-L1 mRNA (blue) and PD-L1-lnc (red) in A549 and PC9 cells. **(E)** The expression level of PD-L1 mRNA and PD-L1-lnc in various lung adenocarcinoma cancer cell lines detected by qRT-PCR with specific primers.

To further investigate whether the PD-L1-lnc exist in other cancers in addition to LUAD, we analyzed the RNA-seq data in TCGA database. The results showed the expression of PD-L1-lnc in various cancers, including BRAC, ESCA and STAD, etc. (fig. S4A). Pan-cancer analysis in TCGA database further indicated a negative association of PD-L1-lnc level with cancer patient survival rates (fig. S4B). To confirm these results derived from TCGA database, we used specific probes to detect both PD-L1 mRNA and PD-L1-lnc in multiple cancer cell lines and cancer tissues (fig. S4C, left). The PCR products confirmed the expression of PD-L1-lnc in various tumor cells and tissues (fig. S4C, right).

We next linked PD-L1-lnc to GFP mRNA in an expressing system to confirm PD-L1-lnc as a non-coding RNA (fig. S5A, upper). RT-PCR assay and fluorescence microscopy showed a high level of GFP mRNA (fig. S5A, lower) but no GFP protein in the PD-L1-lnc-expressing system (fig. S5B), confirming PD-L1-lnc as a lncRNA.

### Lung cancer cell PD-L1-lnc is markedly upregulated by IFNγ in a similar manner to PD-L1 mRNA

The expression of PD-L1 mRNA and protein in tumor cells can be upregulated by T cell-secreted IFNγ [24], which provides a mechanism for cancer to suppress T cell immune responses via ligating T cell surface PD-1. Given that PD-L1-lnc is spliced from 638 to 744 and 832 to 899 nucleotides downstream of the splicing site of PD-L1 mRNA transcription (Fig. 3A), we anticipated that IFNγ would enhance PD-L1-lnc transcription, in a similar manner to that of PD-L1 mRNA. To test this, we treated A549, PC9, H1975, H1650 and H1299 cells with or without IFNγ followed by detection of PD-L1 protein, PD-L1 mRNA and PD-L1-lnc levels. Western blot analysis showed that IFNγ treatment strongly enhanced PD-L1 protein level in the cells (fig. S6A). Increase of PD-L1 protein level on cancer cell surface was also shown by flow cytometry analysis (fig. S6B). In agreement with previous report [25], IFNγ treatment strongly increased PD-L1 mRNA level in A549, PC9, H1975, H1650 and H1299 cells (Fig. 3B). In a similar manner, IFNγ treatment also significantly increased PD-L1-lnc level in all the lung cancer cell lines (Fig. 3B). To further examine the distribution of PD-L1-lnc in lung cancer cells, we purified nuclear and cytoplasm fractions from A549 cells. As shown in Fig. 3C, left, isolated nuclear and cytoplasm fractions exhibited high levels of marker proteins, Histone H3 and GAPDH, respectively, confirming the enrichment of nuclear and cytoplasm fraction in isolated products. Agarose gel (Fig. 3C, left) and RT-PCR analysis (Fig. 3C, right) demonstrated that PD-L1-lnc and PD-L1 mRNA in both nuclear and cytoplasm fractions were upregulated by IFNγ treatment.

**Figure 3.**
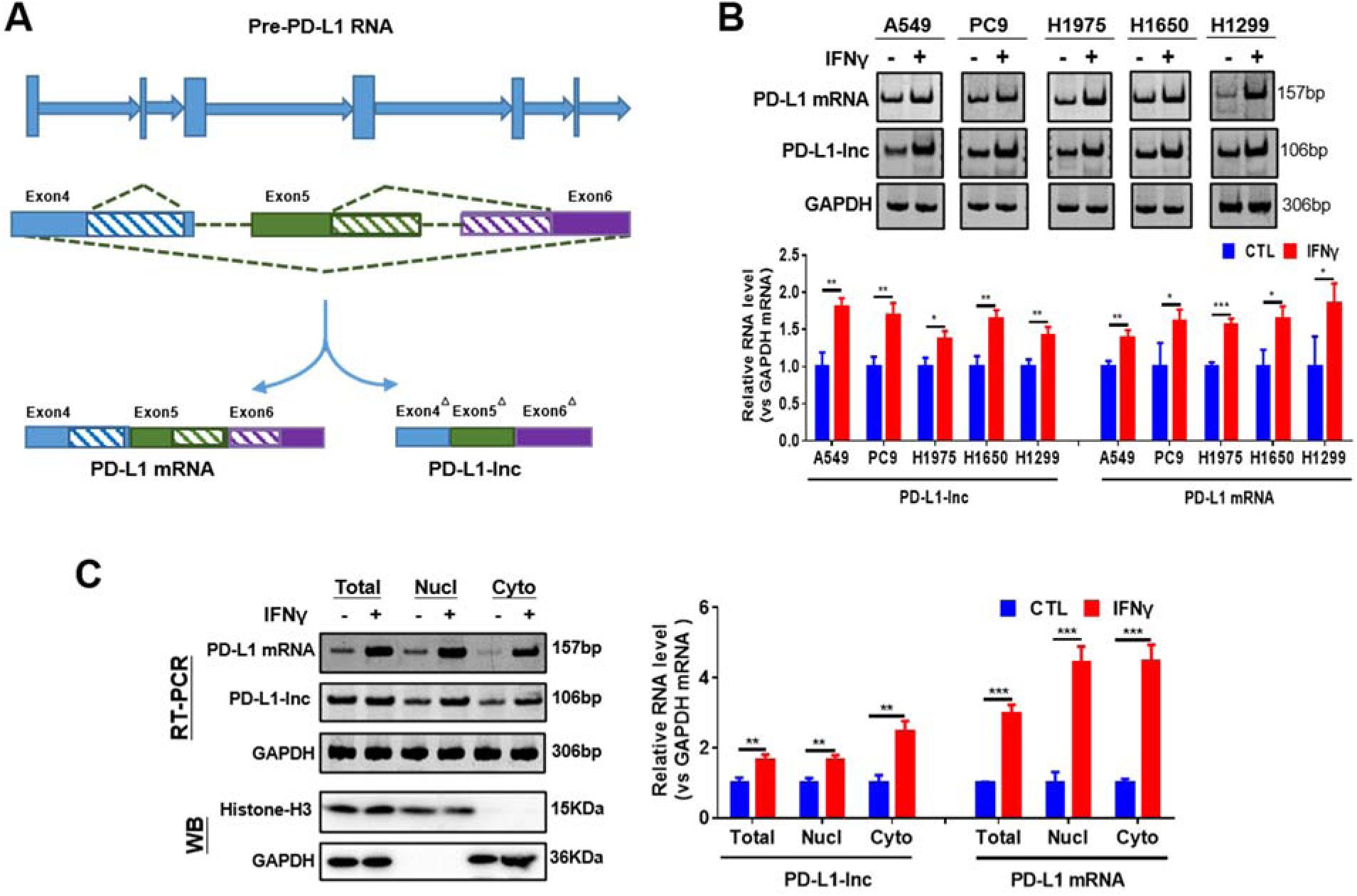
Human lung adenocarcinoma PD-L1-lnc was regulated by IFNγ treatment. **(A)** Schematic representation of the alternative splice region of the PD-L1 variants. **(B)** Agarose gel analysis of the expression level of PD-L1 mRNA and lncRNA in lung adenocarcinoma cells with or without IFNγ treatment. Upper: representative image; Lower: quantitative analysis. **(C)** Cellular distribution of PD-L1-lnc and PD-L1 mRNA in A549 cells with or without IFNγ treatment. Left: representative image; right: quantitative analysis. * *P* < 0.05; \*\**P* < 0.01; \*\*\**P* < 0.001.

### PD-L1-lnc promotes proliferation and invasion but suppresses apoptosis of lung cancer cells

It has been reported that lncRNAs derived from gene alternative splicing can protect the corresponding gene mRNA against the nonsense-mediated decay (NMD) pathway, which targets mRNAs harboring premature termination codons for degradation [26]. Since PD-L1-lnc use the same 5’-most supported translational start codon as the PD-L1 mRNA, the PD-L1-lnc was rendered a candidate for nonsense-mediated mRNA decay to protect PD-L1 mRNA. To test this, we firstly compared the expression level of PD-L1-lnc and PD-L1 mRNA in our collected lung cancer tissues and lung adenocarcinoma data set from TCGA. The results showed there was no correction between PD-L1-lnc and PD-L1 mRNA in both our collected lung cancer tissues and lung adenocarcinoma data set from TCGA (fig. S7A-B). Moreover, we also analyzed associations between PD-L1-lnc or PD-L1 mRNA and overall survival of lung adenocarcinoma based on the TCGA database. Notably, the Kaplan–Meier survival analysis showed that lung adenocarcinoma patients with high PD-L1-lnc expression had shorter overall survival (fig. S7C), while the expression level of PD-L1 mRNA had no effect on the overall survival of lung adenocarcinoma patients (fig. S7D). In order to further explore the relationship between PD-L1-lnc or PD-L1 mRNA, we overexpressed PD-L1-lnc by PD-L1-lnc vector or depleted PD-L1-lnc by PD-L1-lnc shRNA in lung cancer cells and then monitored the cellular levels of PD-L1 mRNA and protein. As showed in fig. S7E, the expression level of PD-L1 mRNA had no change despite PD-L1-lnc was significantly upregulated or downregulated. Western blotting and Flow cytometry analysis also showed overexpression or depletion of PD-L1-lnc in A549 cells had no effect on the protein level of PD-L1 (fig. S7F-G). We further monitored the effect of PD-L1-lnc on the stability of PD-L1 mRNAs, the decay of PD-L1 mRNAs in A549 cells transfected with control vector and PD-L1-lnc vector was assessed by qRT-PCR after blocking the transcription of new PD-L1 mRNAs via actinomycin D (ActD) treatment. As shown in fig. S6H, the relative level of PD-L1 mRNAs in A549 cells transfected with control vector and PD-L1-lnc vector displayed no difference. Furthermore, the effect of PD-L1-lnc on the expression of PD-L1 mRNA and protein during IFNγ stimulation was also investigated. Both flow cytometry and Western blots analysis showed that PD-L1-lnc had no effect on the PD-L1 protein level in A549 cells before or after IFNγ treatment (fig. S7I-J). Taken together, inconsistent with our assumptions and the prediction by NCBI database, PD-L1-lnc exhibited no effect on PD-L1 mRNA expression.

Although PD-L1-lnc could not enhance the expression of PD-L1 mRNA and protein, we found that it strongly affected lung cancer progression. In these experiments, cancer cell proliferation, invasion and apoptosis were assessed using EdU staining [27], Transwell assay [28] and Annexin V labeling [29], respectively. As shown in Fig. 4, overexpression of PD-L1-lnc in A549 and PC9 lung cancer cells strongly enhanced cell proliferation (Fig. 4A) and invasion (Fig. 4B) but suppressed tumor cell apoptosis (Fig. 4C) compared to untreated cancer cells. In contrast, when PD-L1-lnc in lung cancer cells was depleted via transfection with specific PD-L1-lnc shRNA, the proliferation (Fig. 4A) and invasion (Fig. 4B) of cancer cells were significantly reduced, whereas the apoptosis of cancer cells (Fig. 4C) was increased, compared to untreated cancer cells.

**Figure 4.**
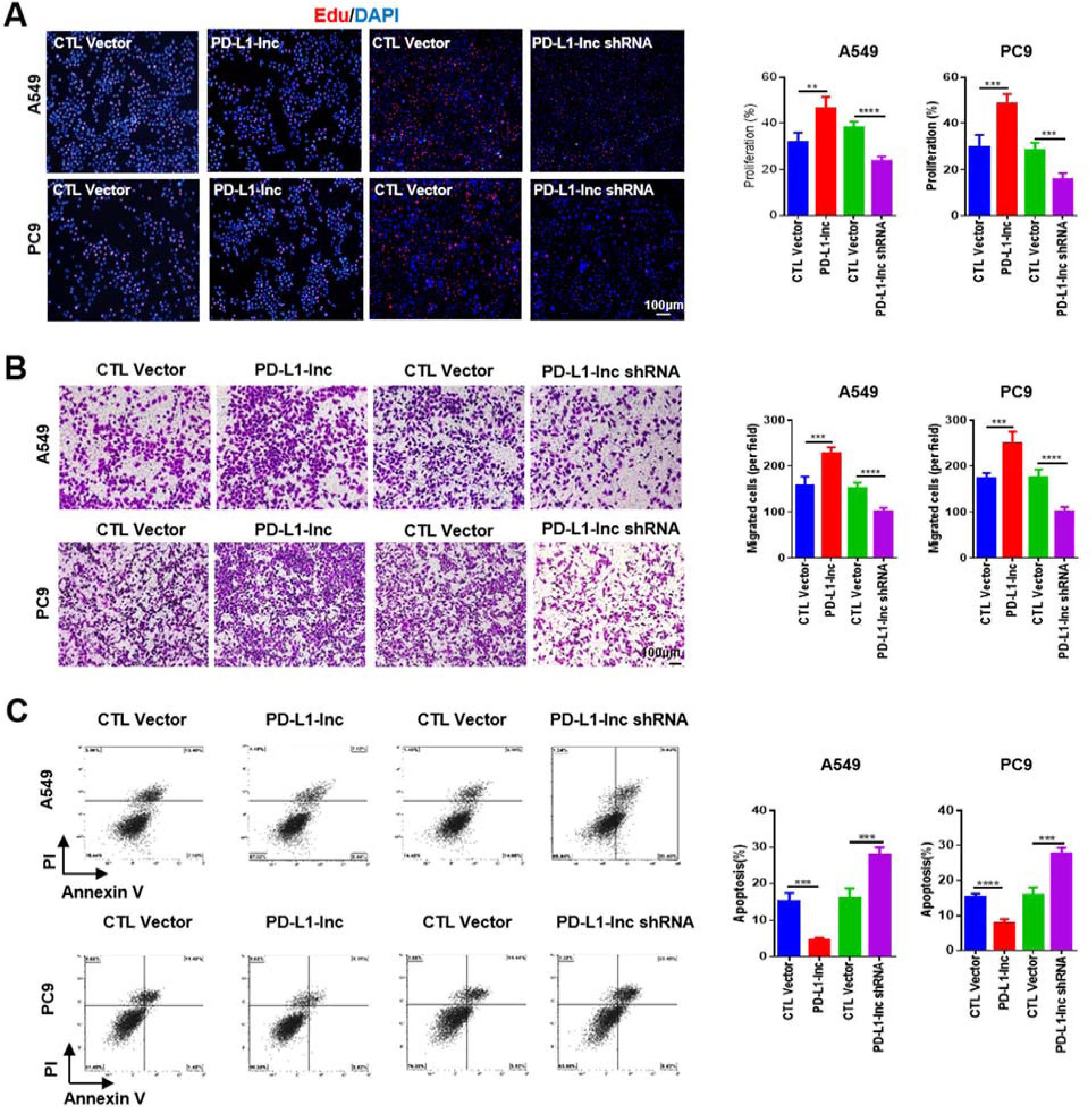
Effects of PD-L1-lnc on the proliferation, invasion and apoptosis of human lung cancer cells. A549 and PC9 cells were transfected with PD-L1-lnc-expressing vector or PD-L1-lnc shRNA-expressing vector. **(A)** The proliferation of A549 and PC9 cells detected by Edu assay. **(B)** The invasion of A549 and PC9 cells detected by Transwell assay. **(C)** The apoptosis of A549 and PC9 cells detected by flow cytometry. In a-c, left: representative image; right: quantitative analysis. \*\**P* < 0.01; \*\*\**P* < 0.001; **** *P* < 0.0001.

Effects of PD-L1-lnc on the progression of lung cancer cells were further validated in lung cancer xenograft mouse model. For this experiment, we developed two lung cancer cell lines: A549 cells which stably express high level of PD-L1-lnc (PD-L1-lnc) and A549 cells with PD-L1-lnc depleted (PD-L1-lnc shRNA). These two lines of cancer cells, as well as A549 cells transfected with control vector as the negative control, were subcutaneously injected into athymic nude mice. Although no significant difference in body weight was observed in three groups of mice during the time frame of experiment (Fig. 5A), tumor size measurement showed a markedly different growth rate among these lung cancer cells (Fig. 5B-C). Specifically, A549 cells expressing higher level of PD-L1-lnc grew significantly faster than control A549 cells, whereas A549 cells with PD-L1-lnc depletion showed a growth delay compared to control A549 cells. In line with this, Ki67 staining analysis in tumor tissue sections showed that A549 cells expressing higher level of PD-L1-lnc had a higher proliferation rate whereas A549 cells with PD-L1-lnc depletion had a lower proliferation rate than control A549 cells (Fig. 5D). Analysis of PD-L1-lnc level in the implanted tumor tissues by RT-PCR confirmed that, compared to control A549 lung cancer tissue, PD-L1-lnc A549 tumor and PD-L1-lnc shRNAA549 tumor expressed significantly higher and lower level of PD-L1-lnc, respectively (Fig. 5E). In contrast, PD-L1 mRNA levels in both PD-L1-lnc and PD-L1-lnc shRNA A549 tumors were not different.

**Figure 5.**
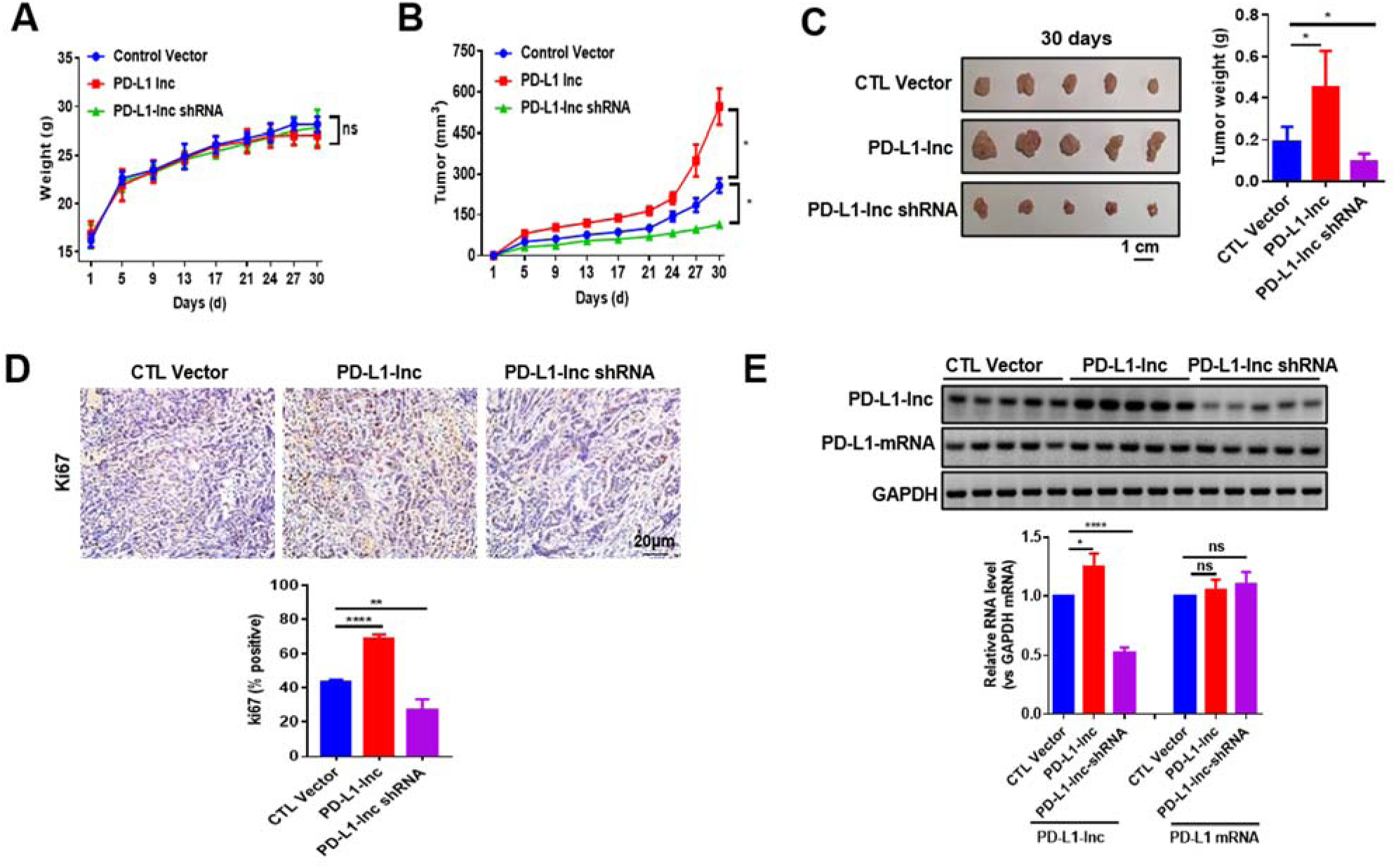
Effects of PD-L1-lnc on the growth of lung cancer cell xenografts in mice. **(A)** The growth rate of mouse weight during the timeframe of experiment. **(B)** The growth rate of various tumor cells implanted in the mice. **(C)** The weight of various implanted tumors on 30 days post-injection. **(D)** Proliferation of implanted tumors in the mice. Upper: representative immunohistochemical staining of Ki67 in implanted tumors; Lower: quantitative analysis. **(E)** qRT-PCR analysis of the expression levels of PD-L1-lnc in the implanted tumors. Upper: gel image of PCR product; Lower: qRT-PCR analysis. ns, no significance. \**P* < 0.05; \*\**P* < 0.01.

### PD-L1-lnc executes its oncogenic function through activating c-Myc signaling

To explore the mechanism underlying the pro-tumor function of PD-L1-lnc, we performed microArray analysis of human genes in A549 cells which stably transfected with PD-L1-lnc-expressing vector (PD-L1-lnc) and A549 cells transfected with control vector. As shown in Fig. 6A, we found that 2640 genes were upregulated while 1810 genes were downregulated (fold-change >2) in A549 cells after PD-L1-lnc overexpression. Further analysis of upregulated or downregulated genes compared to control A549 cells indicated that majority genes (∼20.9%) belonged to the c-Myc pathway regulation (Fig. 6B). Next, using qRT-PCR, we examined a panel of these genes that both displayed significant alteration in microArray analysis and were in the c-Myc-regulating network. As shown in Fig. 6C, PD-L1-lnc overexpression or depletion markedly altered these genes in both A549 cells and lung cancer xenograft (Fig. 6C). These results indicated PD-L1-lnc might promote tumor progress through c-Myc.

**Figure 6.**
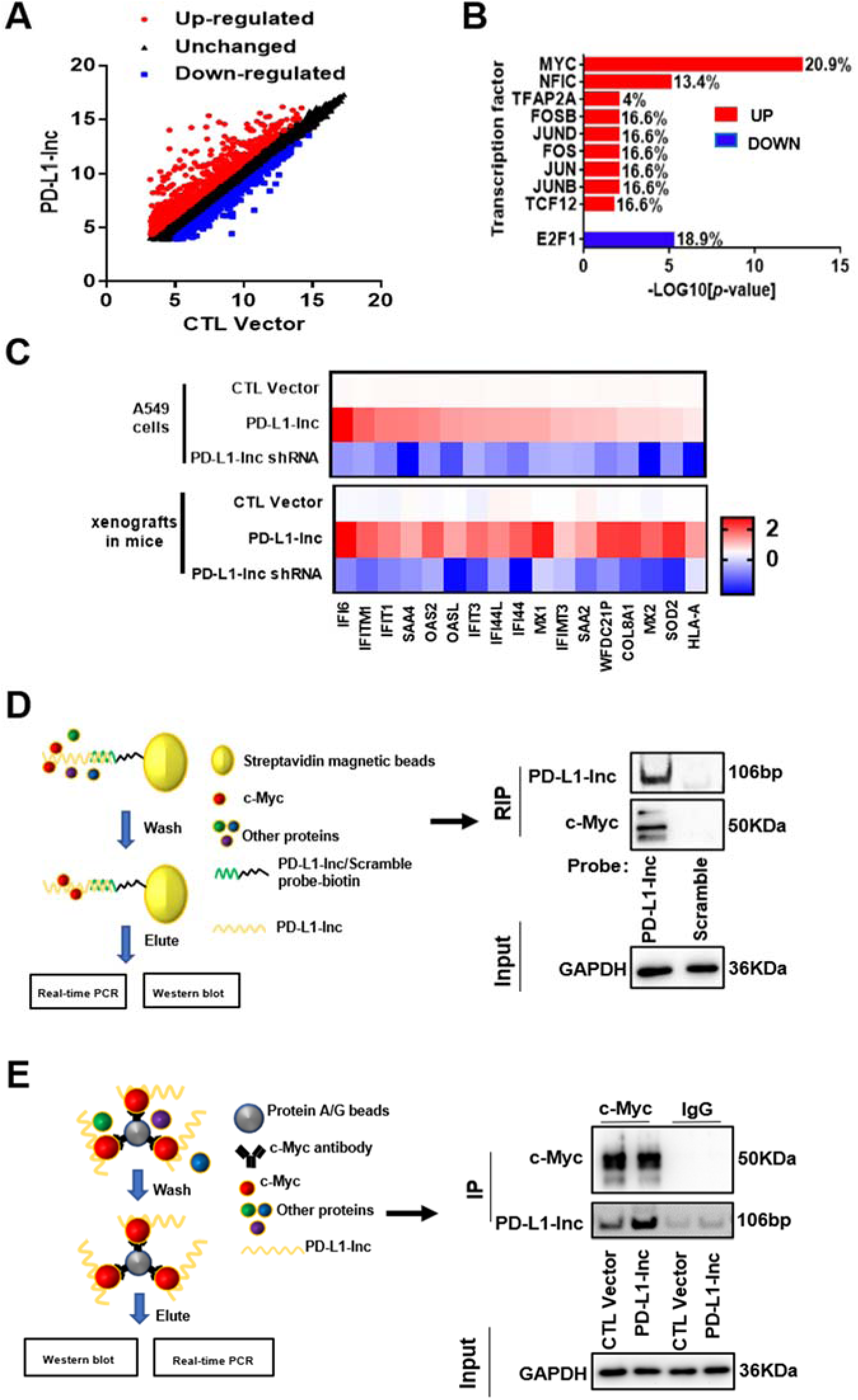
PD-L1-lnc binds to c-Myc and activates c-Myc signal downstream. **(A)** MicroArray analysis of mRNA in A549 cells with PD-L1-lnc overexpression. **(B)** Transcriptional factor analysis among the upregulated or downregulated genes in A549 cells with PD-L1-lnc overexpression. **(C)** Expression level of genes in c-Myc signaling downstream in A549 cells with PD-L1-lnc overexpression or depletion (Upper) and lung cancer xenograft (Lower). **(D)** Pull-down of PD-L1-lnc in A549 cells using biotinylated antisense probes against PD-L1-lnc to detect the PD-L1-lnc-associated c-Myc. **(E)** Pull-down of c-Myc in A549 cells using anti-c-Myc antibody and ProteinA/G beads to detect the c-Myc-associated PD-L1-lnc.

LncRNAs have been confirmed to play important roles in regulating tumorigenesis through interaction with proteins. We used two strategies to examine the potential interactions between PD-L1-lnc and c-Myc, a main member of MYC family that plays a critical role in cancer cell proliferation, metastasis and apoptosis resistance [30]. As shown in Fig. 6D, left, we designed a system consisting of biotin-conjugated antisense oligonucleotide of PD-L1-lnc and streptavidin-conjugated magnetic beads to specifically pull-down PD-L1-lnc. Biotin-conjugated scramble oligonucleotide was used as a negative control. Western blot analysis showed that c-Myc was immunoprecipitated in association with PD-L1-lnc, whereas no c-Myc was detected in immunoprecipitated complex by scramble oligonucleotide (Fig. 6D, right). We also immunoprecipitated c-Myc in A549 cells that were overexpressed with PD-L1-lnc or control A549 cells using anti-c-Myc antibody followed by Protein A/G-conjugated Sepharose beads (Fig. 6E, left). As shown in Fig. 6E, right, PD-L1-lnc was detected in the immunoprecipitated product from the control A549 cells and PD-L1-lnc overexpressed A549 cells by anti-c-Myc antibody, whereas no PD-L1-lnc was detected in immunoprecipitated complex by IgG. Moreover, there was much more PD-L1-lnc in the immunoprecipitated product from PD-L1-lnc overexpressed A549 cells, compared to the control A549 cells. Above all, our results suggested the association of PD-L1-lnc with c-Myc in A549 cells.

The interactions between PD-L1-lnc and c-Myc were next analyzed using catRAPID omics [31] and RPISeq [32] algorithms. The interaction probabilities of PD-L1-lnc and c-Myc predicted by both algorithms were more than 0.90, and the most probable binding area of c-Myc in PD-L1-lnc was located in 500-1000nt of PD-L1-lnc. To explore the PD-L1-lnc tertiary conformation, we first determined its secondary structures using minimum free energy algorithm implemented in Mfold (version 2.3) [33]. The results showed that PD-L1-lnc might form hairpin structure, likely from 500nt to 1000nt. This secondary structure with a lower theoretical value of free energy was then selected as a model for 3D structure prediction. The output file containing primary sequence and an associated secondary structure (Dot-Bracket Notation) was submitted to RNA Composer to generate the 3D structure [34]. NPDock [35] was then used to construct the in-silico molecular docking between PD-L1-lnc and c-Myc. The c-Myc 3D structure used in the docking procedure was derived from Protein Data Bank. As showed in Fig. 7A, c-Myc could bind to the double helix structure of hairpin structure formed by PD-L1-lnc. Further analysis showed that hydrogen bonds might be formed between the ASN^100^ of c-Myc and the AG^901-902^ of PD-L1-lnc, the GLY^103^ of c-Myc and the U^883^ of PD-L1-lnc, the PHE^107^ of c-Myc and the G^882^ of PD-L1-lnc, the ALA^110^ of c-Myc and the U^881^ of PD-L1-lnc, the THR^117^ of c-Myc and the AU^880-881^ of PD-L1-lnc, the GLU^118^ of c-Myc and the G^902^ of PD-L1-lnc, the GLY^121^ of c-Myc and the G^877^ of PD-L1-lnc, respectively (fig. S8). These hydrogen bonds could enable the formation of a basic helix-loop-helix Zip motif [36] in c-Myc. To validate these predicted binding sites, two PD-L1-lnc vectors, in which the predicted binding sites were mutated (PD-L1-lnc mut) or depleted (PD-L1-lnc del), were constructed (Fig. 7B). These mutant PD-L1-lnc vectors as well as the WT PD-L1-lnc vector were then transfected into A549 cells. As shown in fig. S9A-B, both WT PD-L1-lnc, PD-L1-lnc del and PD-L1-lnc mut vectors expressed PD-L1-lnc along with its linked GFP mRNA without affecting PD-L1 mRNA and protein. Immunoprecipitation assay using biotin-conjugated antisense oligonucleotide of GFP mRNA plus streptavidin-conjugated magnetic beads showed that c-Myc was associated with wild type PD-L1-lnc but not mutant PD-L1-lnc (Fig. 7C). In line with this, direct pulldown of c-Myc using anti-c-Myc antibody plus Protein A/G-conjugated Sepharose beads in A549 cells transfected with WT PD-L1-lnc-GFP mRNA or mutant PD-L1-lnc-GFP mRNA indicated that only WT PD-L1-lnc-GFP mRNA but not mutant PD-L1-lnc-GFP mRNA was associated with c-Myc (Fig. 7D).

**Figure 7.**
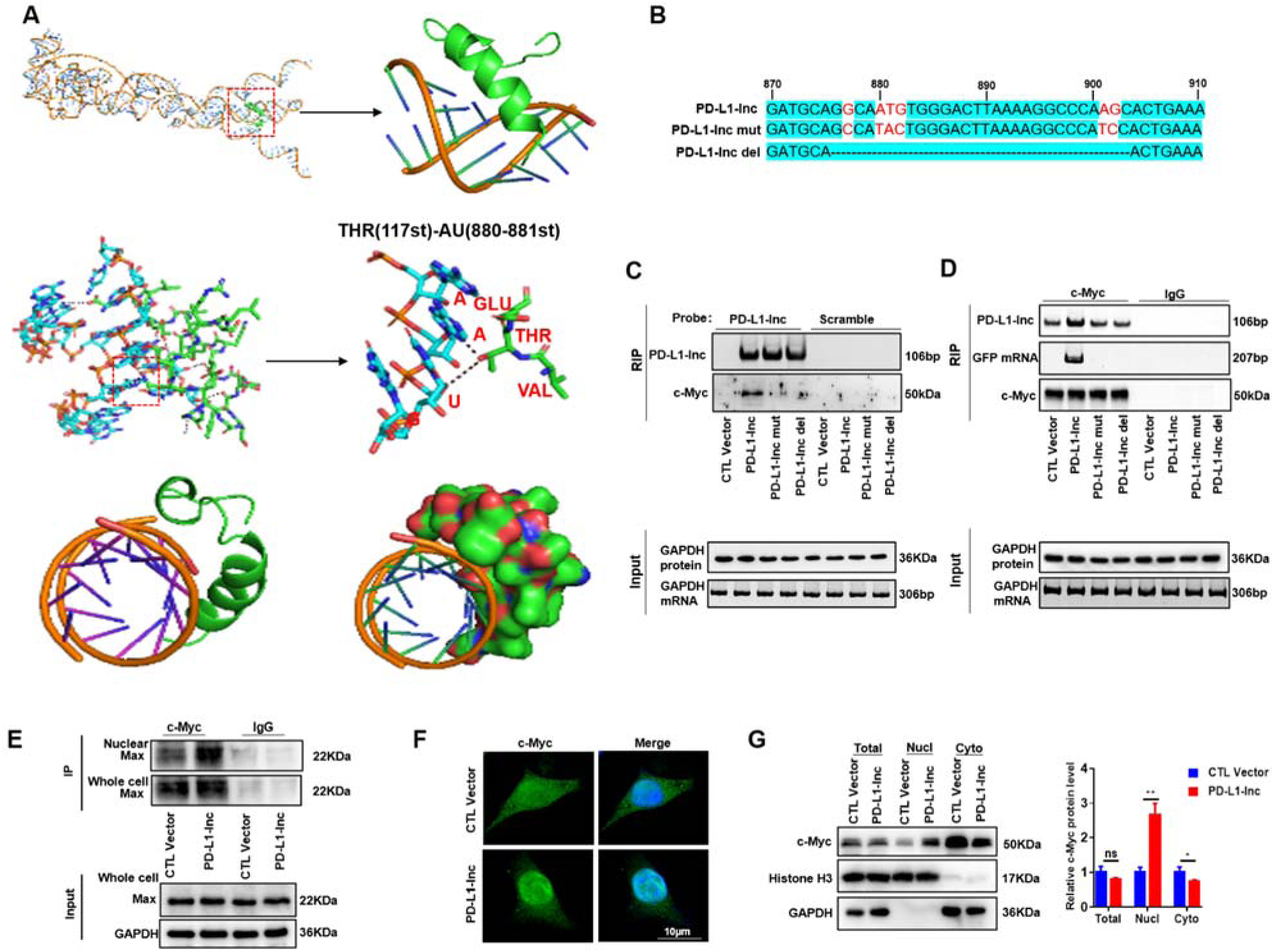
Binding of PD-L1-lnc with c-Myc enhances the association of c-Myc with MAX and promotes the entry of c-Myc into nucleus. **(A)** Graphical representation of three-dimensional structures of PD-L1-lnc and c-Myc docking models with a zoom-in image of the binding interface. The binding region was shown in three different visualizations (cartoon and sphere). **(B)** Schematic representation of the PD-L1-lnc-del vector which was deleted the predicted binding sites and the PD-L1-lnc-mut vector which was mutated at the predicted binding sites (marked red). **(C)** Pulldown of PD-L1-lnc in A549 cells transfected with WT or mutant PD-L1-lnc/GFP vector using biotinylated antisense probes against GFP mRNA to detect the binding of PD-L1-lnc with c-Myc. **(D)** Immunoprecipitation of c-Myc in A549 cells transfected with WT or mutant PD-L1-lnc/GFP vector using anti-c-Myc antibody to assess the binding of PD-L1-lnc with c-Myc. **(E)** Binding of PD-L1-lnc with c-Myc resulted in recruitment of MAX by c-Myc in A549 cells. **(F-G)** Immunofluorescence (F) and cell fractionation/Western blot analysis (G) of c-Myc distribution in A549 cells before and after PD-L1-lnc overexpression. Note that PD-L1-lnc overexpression markedly increases the distribution of c-Myc in the nuclei or nuclear fraction (nucl) of A549 cells.

Previous studies showed that helix-loop-helix Zip motif of c-Myc enables formation of a heterodimer with chaperone protein Max in initiating gene transcription. Given that binding of PD-L1-lnc with c-Myc can facilitate c-Myc to form a basic helix-loop-helix Zip motif (Fig. 7A), we postulate that PD-L1-lnc can enhance c-Myc transcriptional activity through enabling a c-Myc conformation change which allows for a heterodimer formation with Max. To test the hypothesis, we compared the protein level of Max in the immunoprecipitated product by anti-c-Myc antibody in the A549 cells transfected the PD-L1-lnc-expressing vector or control vector. As expected, the protein level of Max significantly increased in the A549 cells transfected with the PD-L1-lnc-expressing vector, compared to the A549 cells transfected with the control vector (Fig. 7E). Immunoprecipitation and Western blot analysis further indicated that overexpression of PD-L1-lnc in A549 cells only enhanced the nuclear distribution of Max which were associated with c-Myc, not the total cellular level of c-Myc-associated Max. Supporting the notion that PD-L1-lnc binds to c-Myc, immunofluorescence labeling showed that overexpression of PD-L1-lnc significantly enhanced nuclear translocation of c-Myc in A549 cells (Fig. 7F). Cell fraction and Western blot analysis indicated the increase of nuclear (Nucl) distribution of c-Myc by overexpression of PD-L1-lnc whereas the total cell level of c-Myc remained unchanged (Fig. 7G). These results suggest that PD-L1-lnc may enhance c-Myc activity by enhancing the binding with MAX and promoting the entry of c-Myc into nuclei.

To further test whether PD-L1-lnc executes its function through c-Myc signaling pathway, we knocked down c-Myc expression via c-Myc siRNA in A549 cells (fig. S10) in which PD-L1-lnc was or was not overexpressed and then monitored the cell proliferation, invasion and apoptosis. As shown in Fig. 8A-C, overexpression of PD-L1-lnc strongly promoted A549 cell proliferation (Fig. 8A), invasion (Fig. 8B) and resistance to apoptosis (Fig. 8C). Depleting cellular c-Myc via c-Myc siRNA treatment, however, largely abolished the effect of PD-L1-lnc on enhancing A549 cell proliferation (Fig. 8A), invasion (Fig. 8B) and resistance to apoptosis (Fig. 8C). The function of PD-L1-lnc mutants which do not bind to c-Myc was also investigated. As shown in Fig. 8A-C, transfection with PD-L1-lnc mutants did not enhance cell proliferation, invasion and resistance to apoptosis.

**Figure 8.**
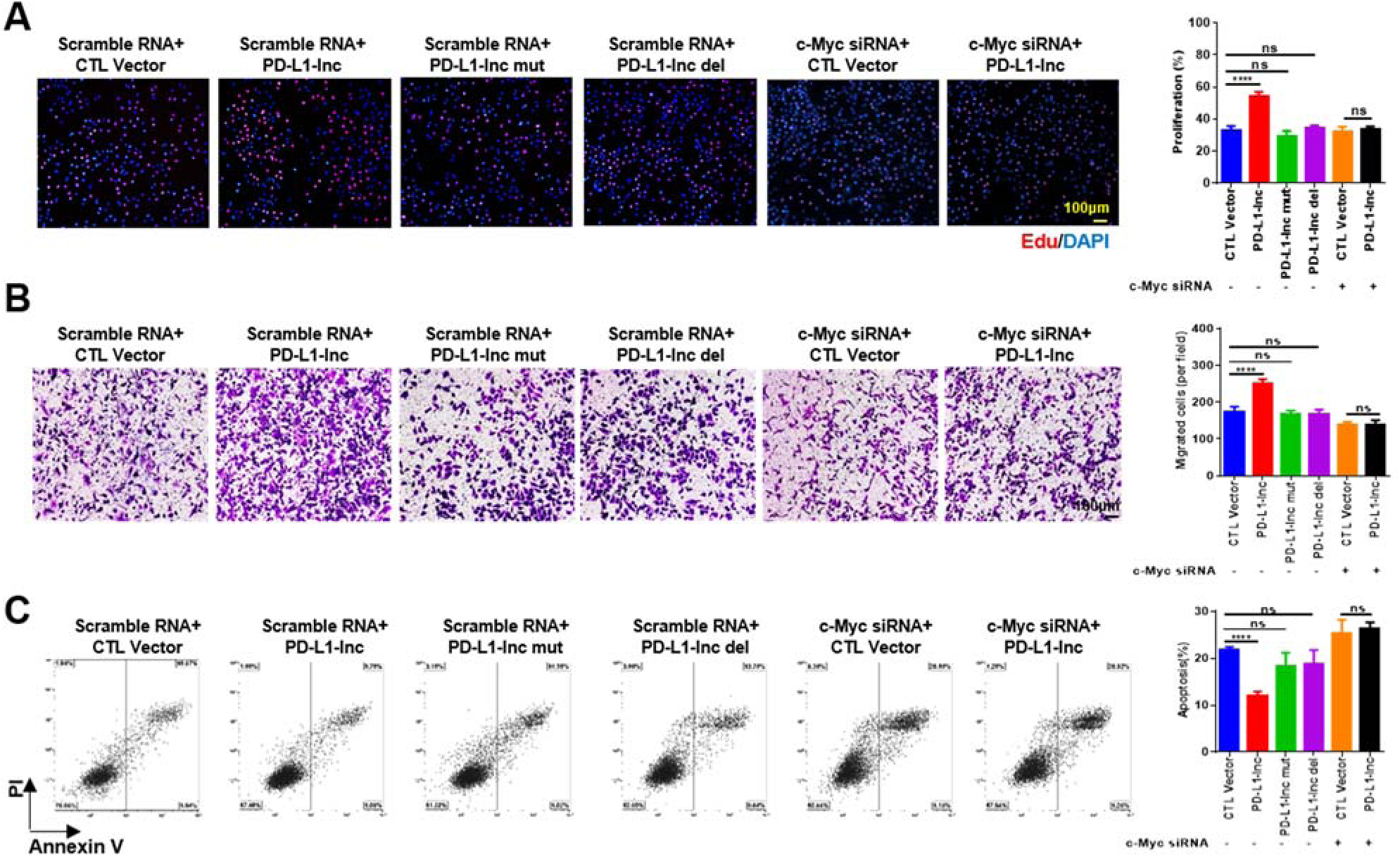
PD-L1-lnc promotes lung adenocarcinoma progression via c-Myc. **(A-C)** Overexpression of PD-L1-lnc mutant or knockdown of c-Myc protein expression in A549 cells both abolished the effect of PD-L1-lnc on A549 cell proliferation (**A**), migration (**B**) and resistance to apoptosis (**C**). In A-C, left panels, representative image from three independent experiments; right panels, quantitative analysis of images. *p < 0.05; **p < 0.01.

## DISCUSSION

As an important immune checkpoint, PD-L1 suppresses T cell immune responses via binding to T cell surface PD-1 and initiating T cell program death [37, 38]. Tumor cells often upregulate PD-L1 expression especially under the condition of T cell infiltration, which provides a mechanism for tumor cells to evade the attack by T cells. Blockade of PD-L1/PD-1 interactions has thus been widely applied in tumor immunotherapy and achieved great success in treating various tumors particularly melanoma, leukemia and lymphoma. However, the efficacy of PD-L1 blockade strategies is limited for solid cancers [6–8]. In the present study, we demonstrate that PD-L1 gene in lung adenocarcinoma tissues, as well as various lung cancer cells, produces another PD-L1 transcript, PD-L1-lnc, via PD-L1 gene alternative splicing. In a similar manner to PD-L1 mRNA, PD-L1-lnc level is markedly increased by IFNγ. Moreover, PD-L1-lnc strongly promotes tumor cell proliferation, survival and invasion via enhancing c-Myc activity. Given that production of PD-L1-lnc in lung cancer cells is not dependent upon PD-L1 protein expression and depletion of PD-L1-lnc markedly suppresses tumor growth both *in vitro* and *in vivo*, depleting tumor cell PD-L1-lnc may provide a novel anti-tumor therapeutic approach in addition to PD-L1 protein blockade.

The efficacy of immune checkpoint blockade in tumor therapy is dependent on the infiltration and activation of anti-tumor T cells in tumor microenvironment, which leads to significant elevation of IFNγ level [39]. Although IFNγ had been regarded as an anti-tumor cytokine [40, 41], recent studies showed a ‘double-edge’ effect of IFNγ in tumor progression [42, 43]. For example, IFNγ can upregulate immune checkpoint molecules particularly PD-L1, which enhances tumor immune evasion [24, 44]. Here we showed that, in addition to PD-L1 mRNA, IFNγ also enhanced expression of PD-L1-lnc. Judged by the PD-L1 gene structure, it seems that PD-L1-lnc and PD-L1 mRNA are upregulated by IFNγ through a similar mechanism.

Previous studies have reported the expression of PD-L1 is modulated by alternative splicing [11, 45]. Zhou et al. identified four PD-L1 splicing variants that lack the transmembrane domain, and these secretory forms of PD-L1 can also suppress the activities of both CD4^+^ and CD8^+^ T cells [46]. In our study, we found that human PD-L1 gene in various lung cancer tissues and cell lines produced a non-coding isoform, NR_052005.1 or PD-L1-lnc, which missed 106nt in exon 4 and 67nt between exon 5 and 6 by alternative splicing (Fig. 3A). Since this isoform lacks an alternate internal segment and uses the 5’-most supported translational start codon as used in mRNA, it may serve as a candidate for nonsense-mediated mRNA decay (NMD) in protecting the mRNA degradation (https://www.ncbi.nlm.nih.gov/gene/29126). However, we found no effect of this isoform on the levels of PD-L1 mRNA and protein, instead, it acts as a functional lncRNA to promote lung adenocarcinoma progress via enhancing c-Myc activity (Fig. 9).

**Figure 9.**
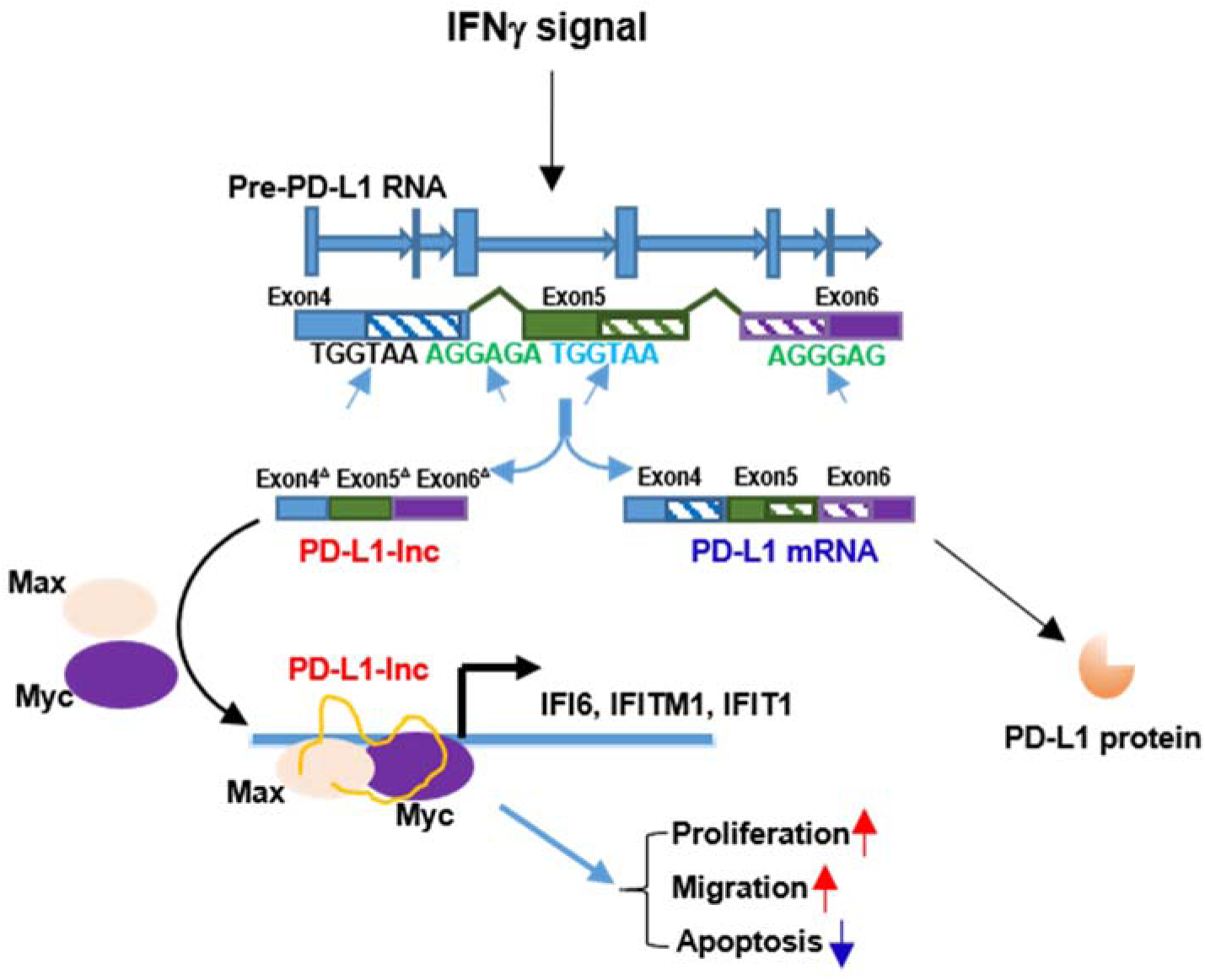
PD-L1 mRNA alternative splice (PD-L1-lnc) promotes lung adenocarcinoma progression via enhancing c-Myc activity.

As a member of transcription factor MYC gene family, c-Myc plays a critical role in the development of human tumors and its overexpression has been detected in lung cancer of different histologic subtypes [47]. Previous study by Tang et al. [48] showed that a long non-coding RNA, LncRNA-GLCC1, could stabilize c-Myc thereby attenuating c-Myc’s ubiquitination with the consequent promotion of colorectal carcinogenesis. Different from their study in which LncRNA-GLCC1 protected c-Myc from ubiquitination and degradation through interacting with HSP90 chaperon, our data suggest that PD-L1-lnc may directly interact with c-Myc. Cross-immunoprecipitation assays clearly showed the association of PD-L1-lnc with c-Myc (Fig. 6D-E). In line with the notion that PD-L1-lnc enhances c-Myc translocation into the nucleus and thus increases its transcriptional activity, we found that PD-L1-lnc overexpression increased the expression of a panel of c-Myc-modulated genes including IFITM1/3, MX1/2, IFI6 and SAA4, etc. Conversely, PD-L1-lnc depletion significantly decreased the expression of these genes.

The interactions between PD-L1-lnc with c-Myc were further validated by bioinformatics analysis using catRAPID omics [31] and RPISeq algorithms [32]. The results suggested that PD-L1-lnc could form hairpin structures, leading to a partial double helix structure. Multiple amino acids of c-Myc could form hydrogen bonds with their corresponding nucleobases in the double helix structure of PD-L1-lnc, which in turn, promotes c-Myc to form a basic helix-loop-helix Zip motif (Fig. 7A), leading to formation of more c-Myc-Max heterodimer and higher c-Myc transcriptional activity.

We have also explored the potential mechanism that modulates alternative splicing for PD-L1-lnc generation. Given that the sequences of splice site in exon 4 and between exon 5 and 6 are UGGUAA, AGGAGA/AGGGAG, respectively. We blasted these sequences in Tomtom database [49] and found that the sequences of splice sites matched to the motifs recognized by three alternative splicing regulators, MSI1, DAZAP1 and ESRP2 (fig. S11A). DAZAP1 was selected to further study since the expression levels of MSI1 and ESRP2 in A549 cells were too low to have function (fig. S11B). To test whether DAZAP1 was involved in the generation of PD-L1-lnc, we checked the expression level of PD-L1-lnc in the A549 cells after transfection the specific siRNAs to DAZAP1. As shown in fig. S11C, the expression level of PD-L1-lnc was significantly decreased (right) after the downregulation of DAZAP1 caused by siRNAs (left), arguing that DAZAP1 may control the generation of PD-L1-lnc. However, further study is required to clarify whether and how DAZAP1 modulate PD-L1-lnc biogenesis.

In summary, the present study reveals that PD-L1-lnc, an alternatively spliced product of the PD-L1 gene, promotes lung adenocarcinoma proliferation, metastasis and survival via enhancing c-Myc transcriptional activity, and provides targeting PD-L1-lnc−c-Myc axis as a novel therapeutic strategy to improve the efficacy of lung cancer therapy.

## Supporting information

manuscripts

## ACCESSION NUMBERS

The data of Microarray analysis in this study can be viewed in GEO database (GSE153051).

## ACKNOWLEGEMENTS

The authors thank Professor Jill Leslie Littrell (Georgia State University, Atlanta, GA) for critical reading and constructive discussion of the manuscript.

## FUNDING

This work was supported by grants from the Ministry of Science and Technology of China (2018YFA0507100), National Nature Science Foundation of China (31801088, 31670917, 31770981), Natural Science Foundation of Jiangsu Province (BK20170076) and Fundamental Research Funds for the Central Universities (020814380095, 020814380082).

## CONFLICT OF INTERESTS

None declared.

## AUTHOR CONTRIBUTIONS

Ken Zen, Hongwei Liang and Tao Wang designed research; Shuang Qu, Zichen Jiao, Geng Lu, Bing Yao, Ting Wang, Weiwei Rong, Jiahan Xu, Ting Fan, Xinlei Sun, and Rong Yang performed research and analyzed data; Jun Wang, Yongzhong Yao, Guifang Xu, Xin Yan, and Tao Wang collected the patients’ samples; Ken Zen and Hongwei Liang wrote the paper.

---

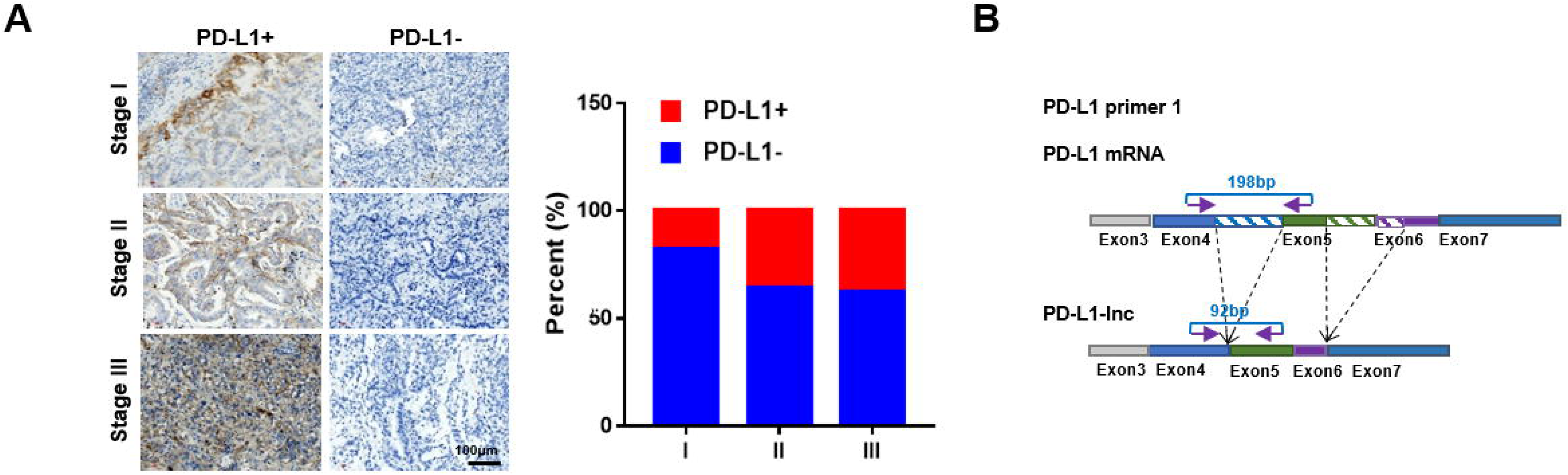

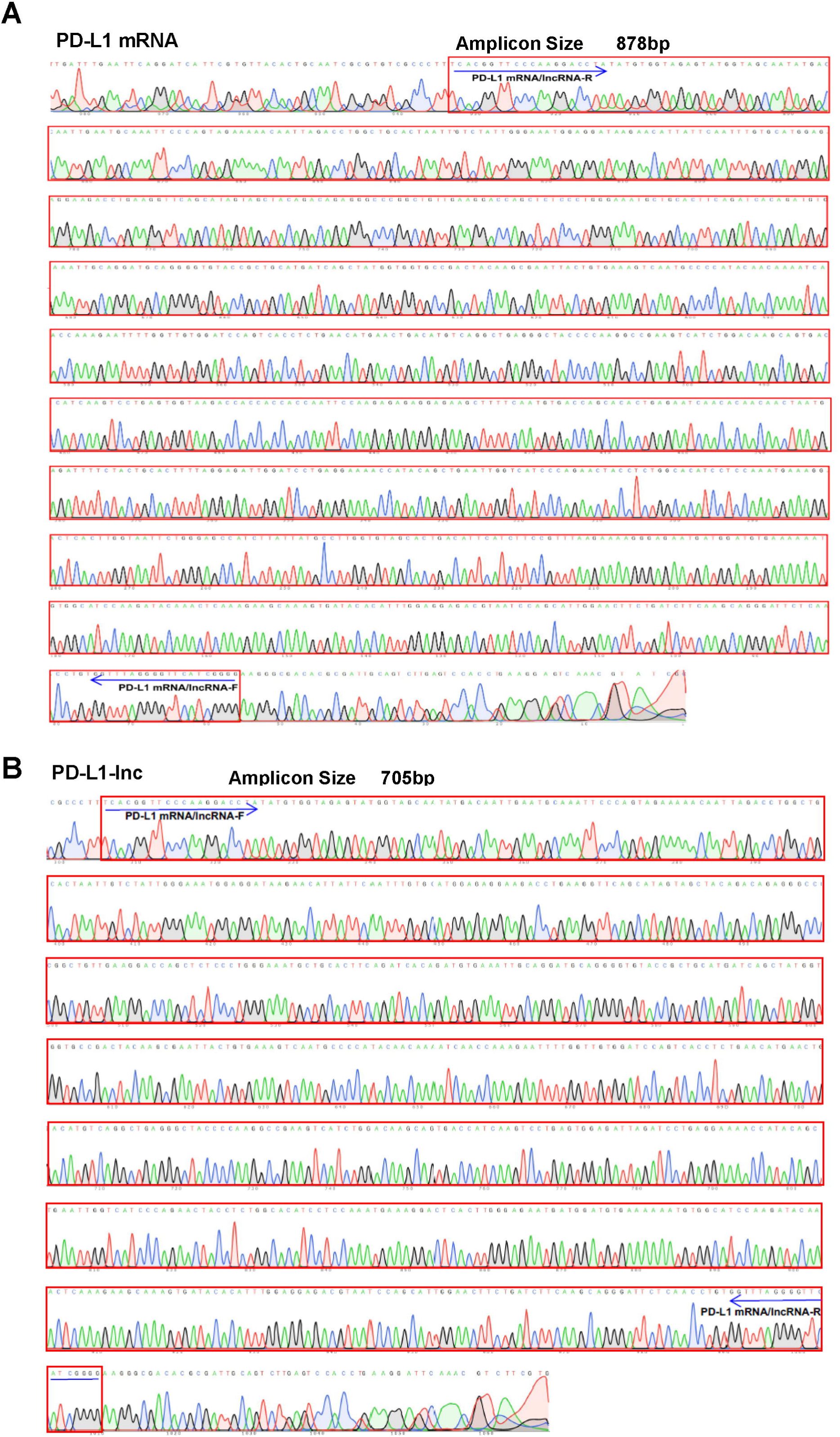

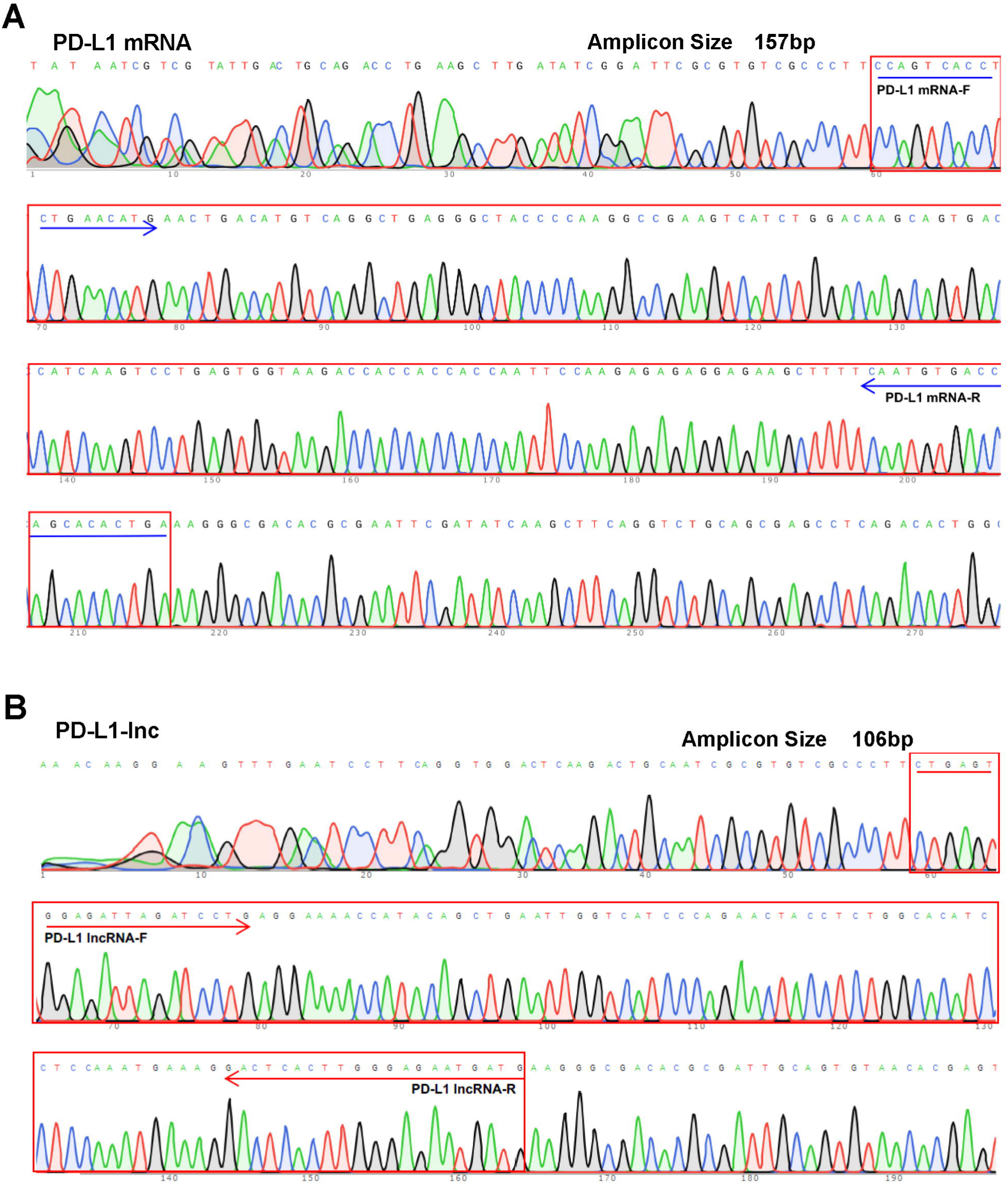

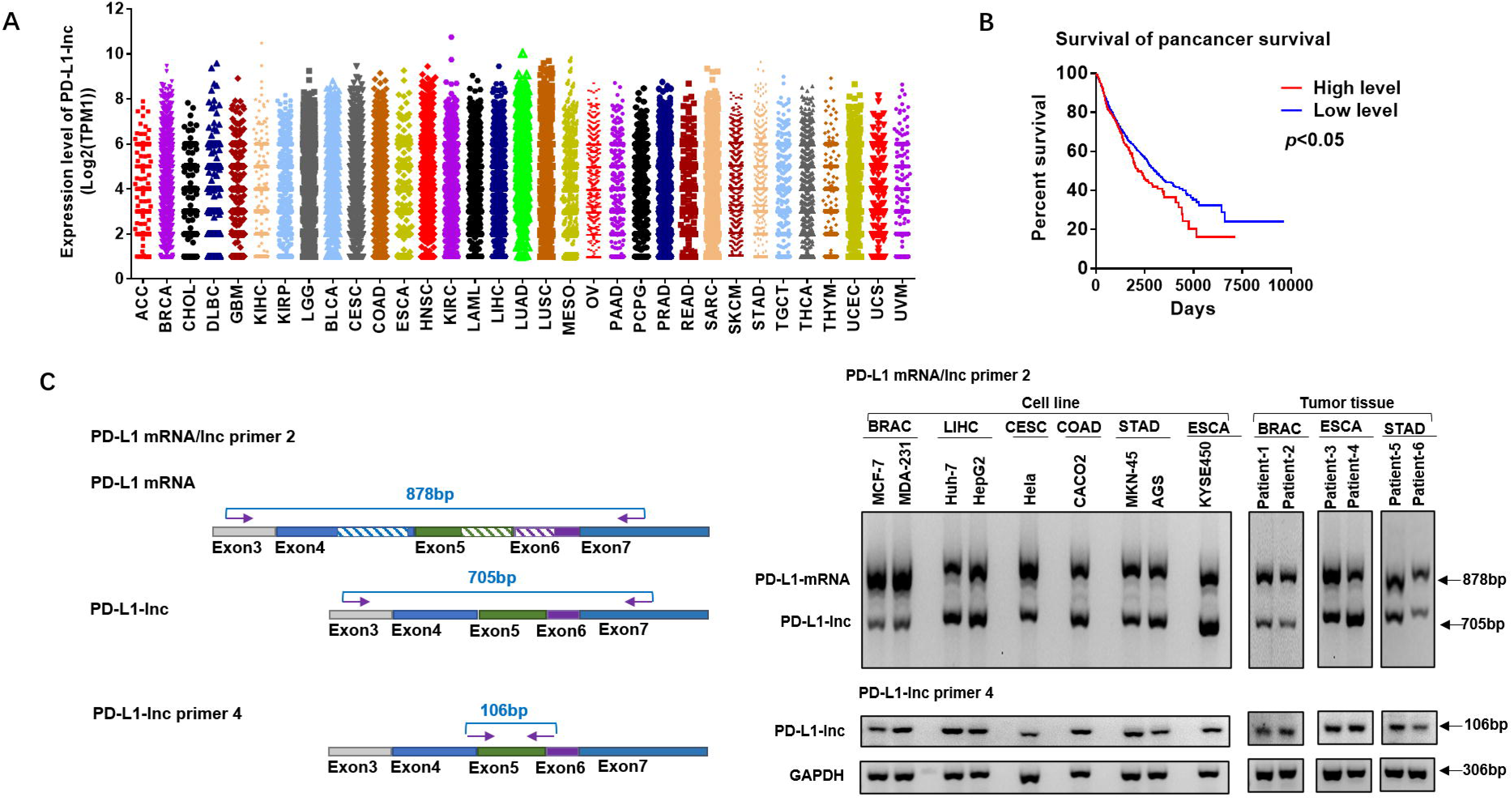

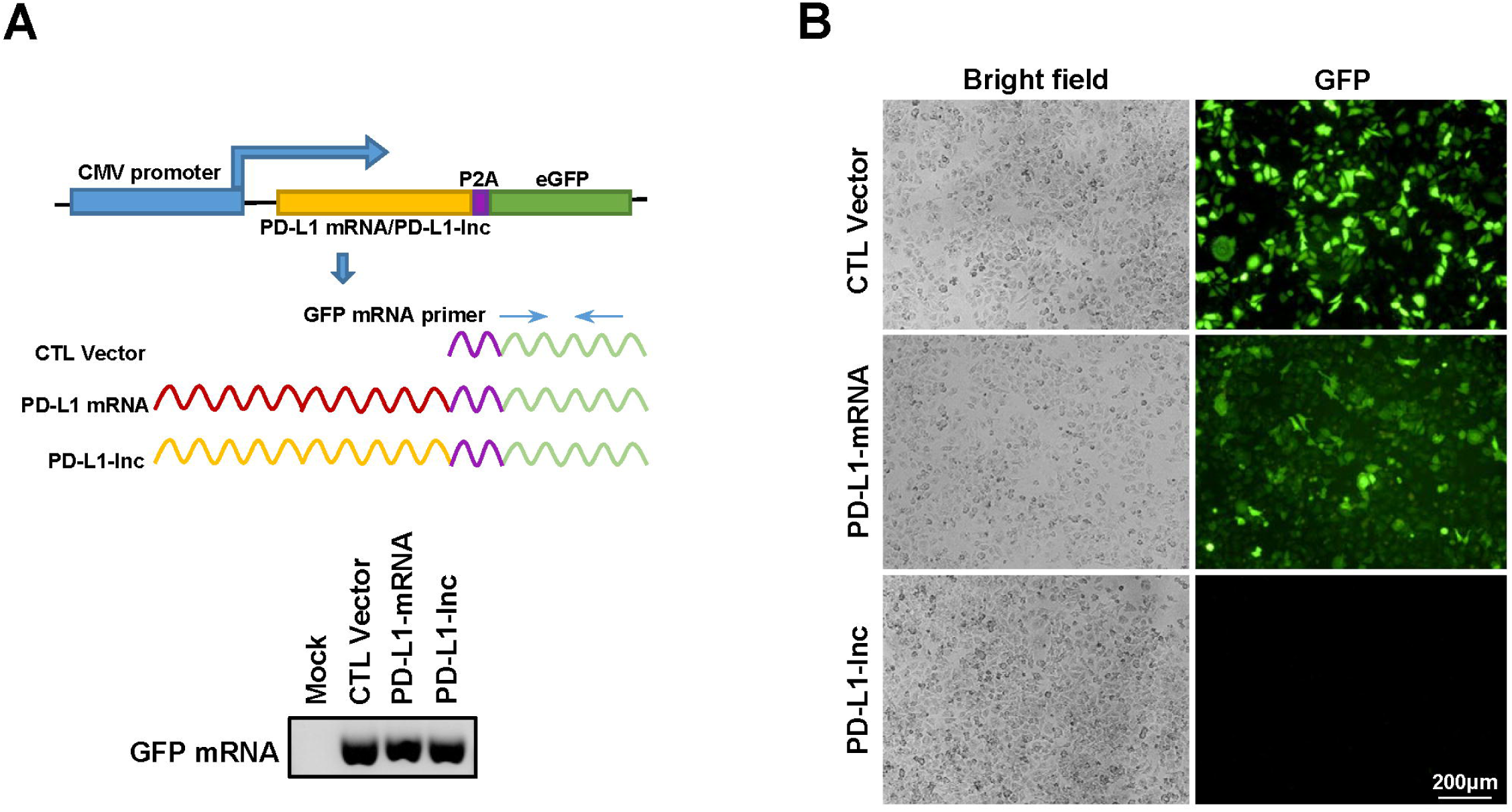

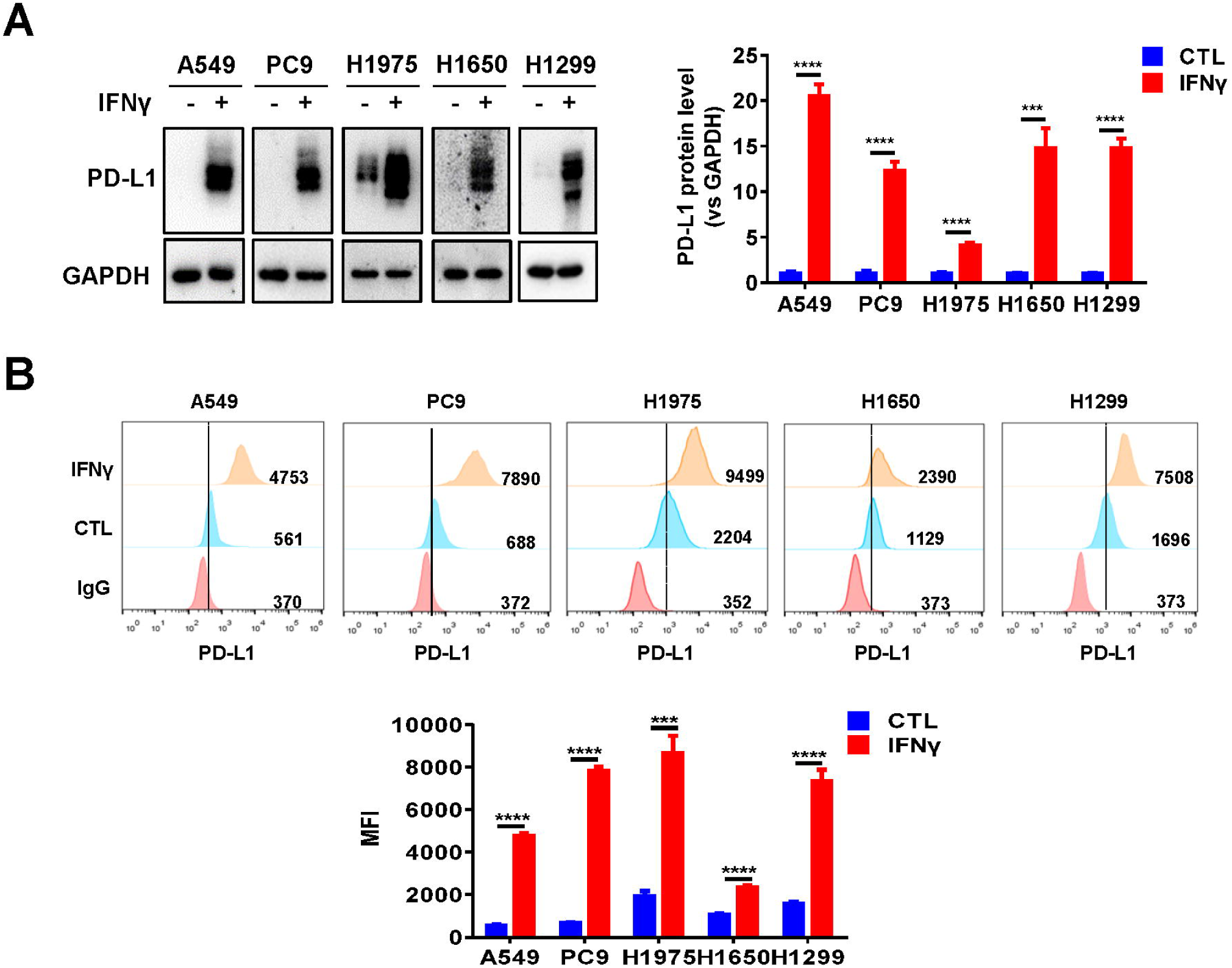

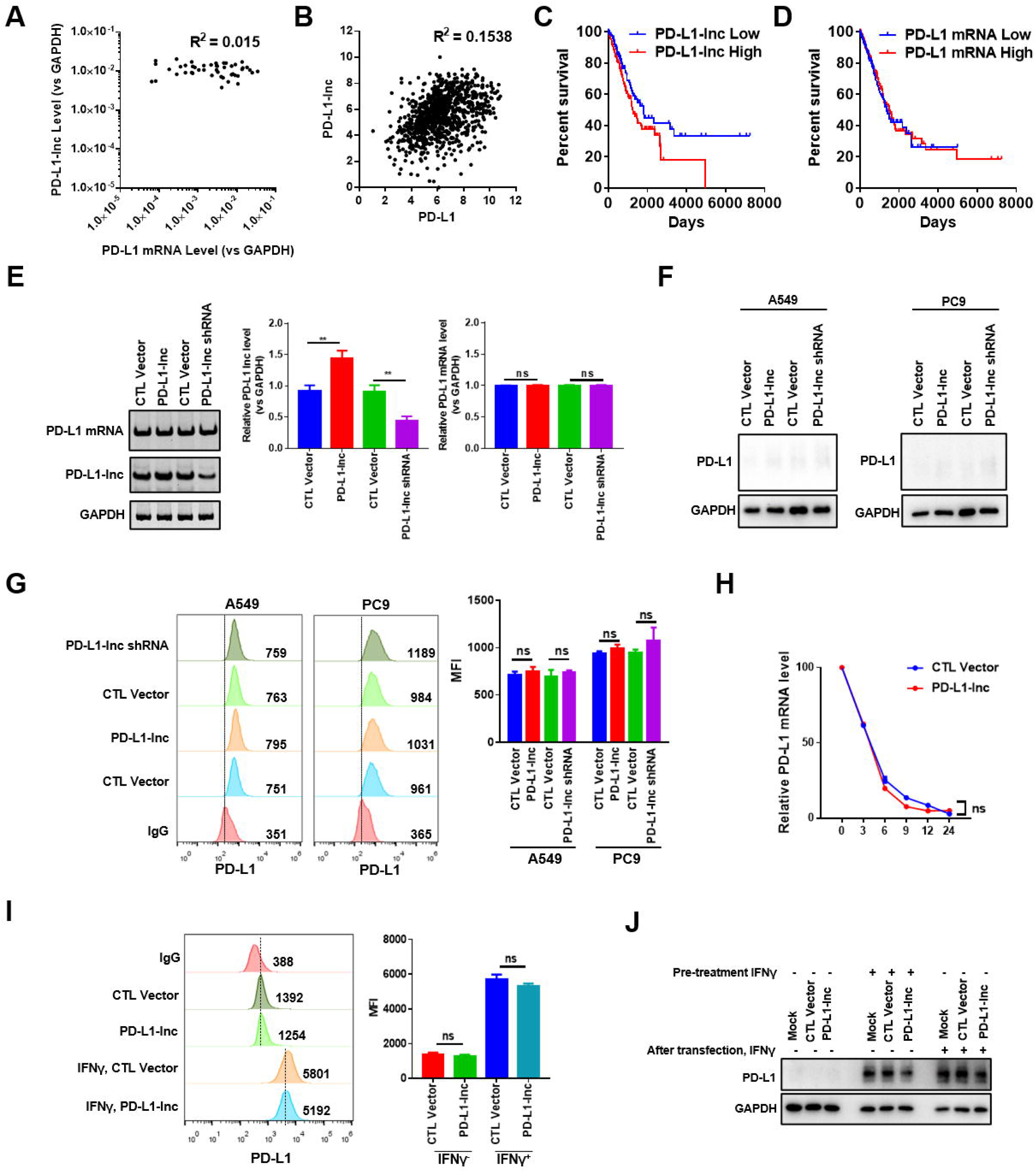

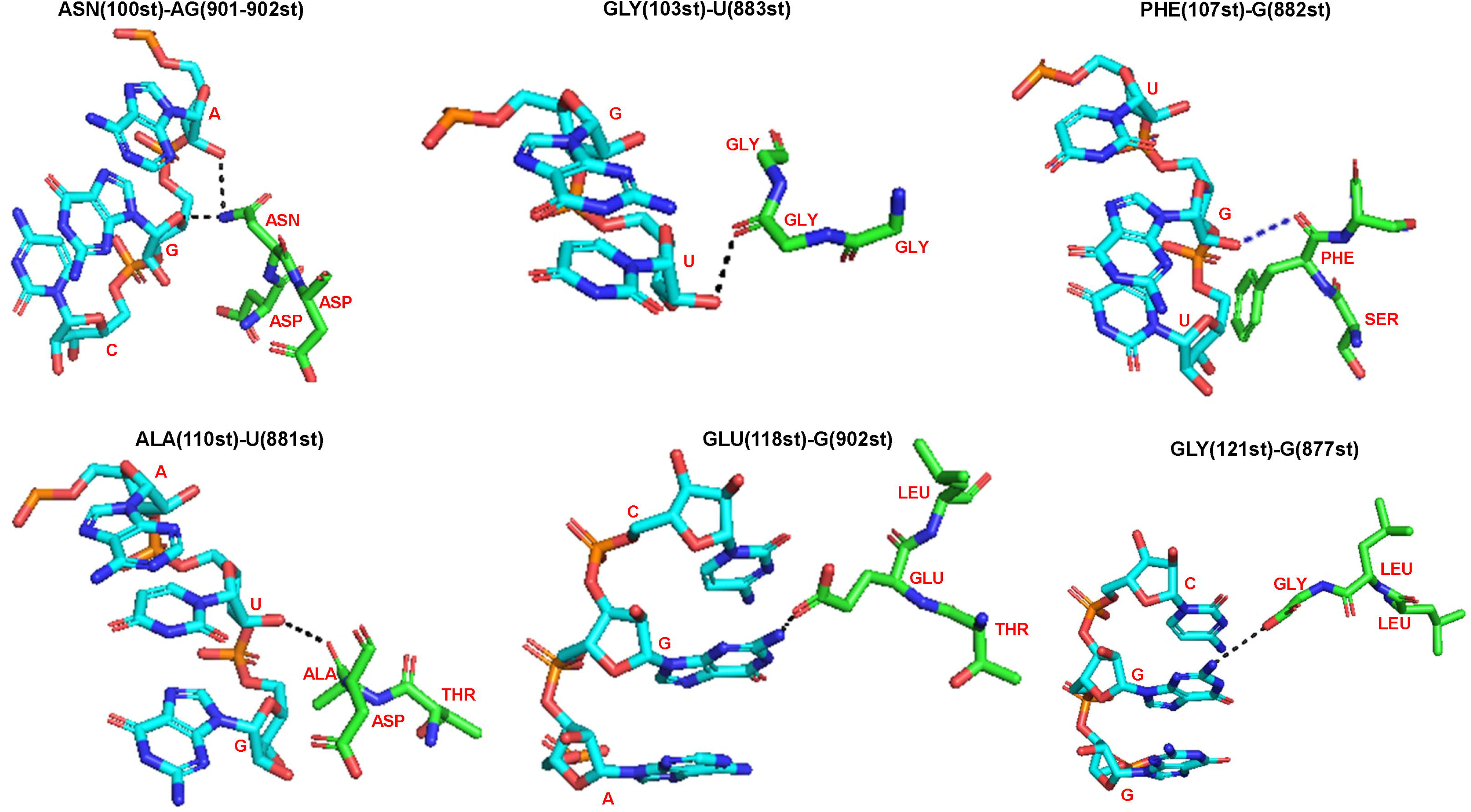

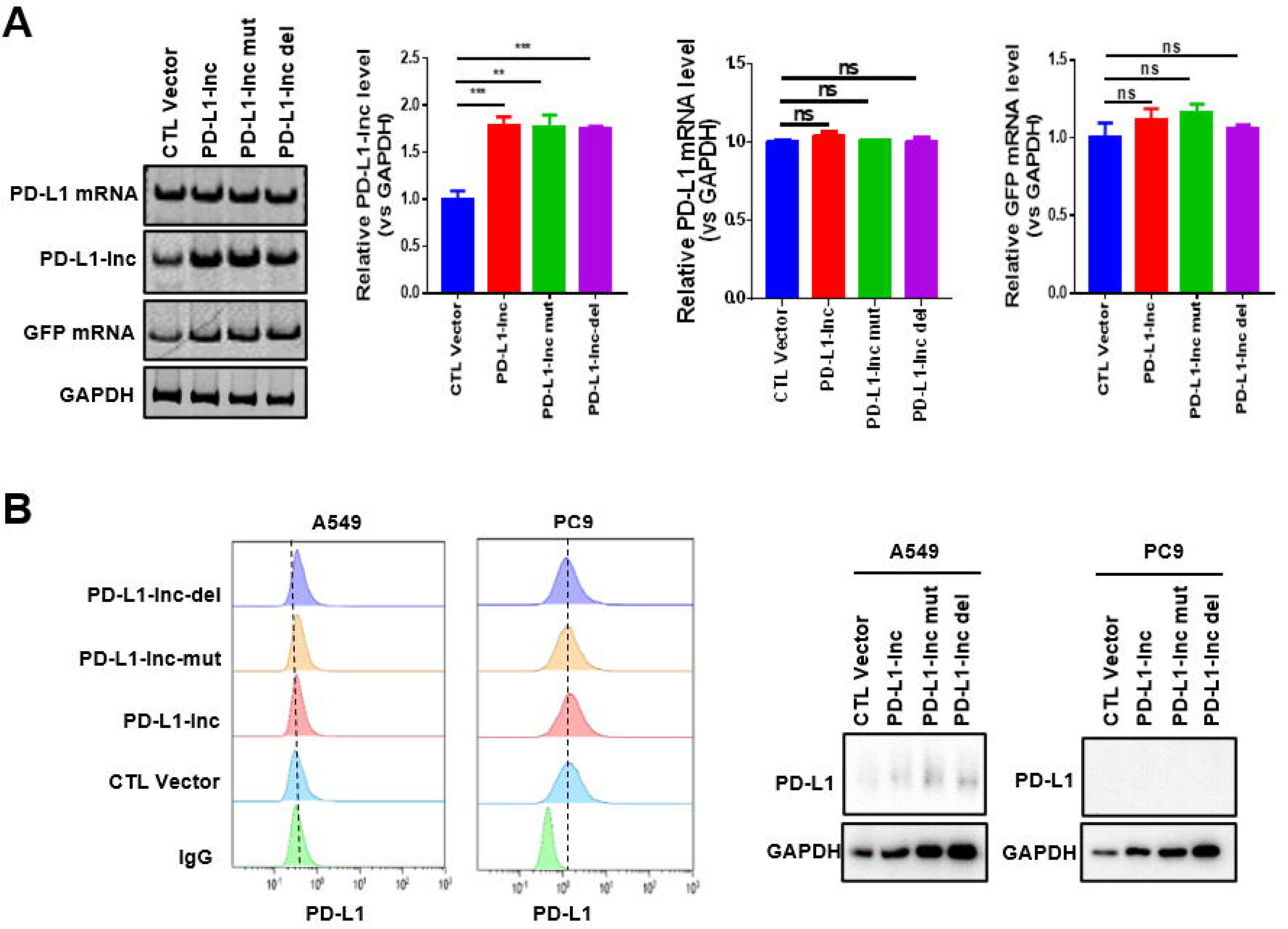

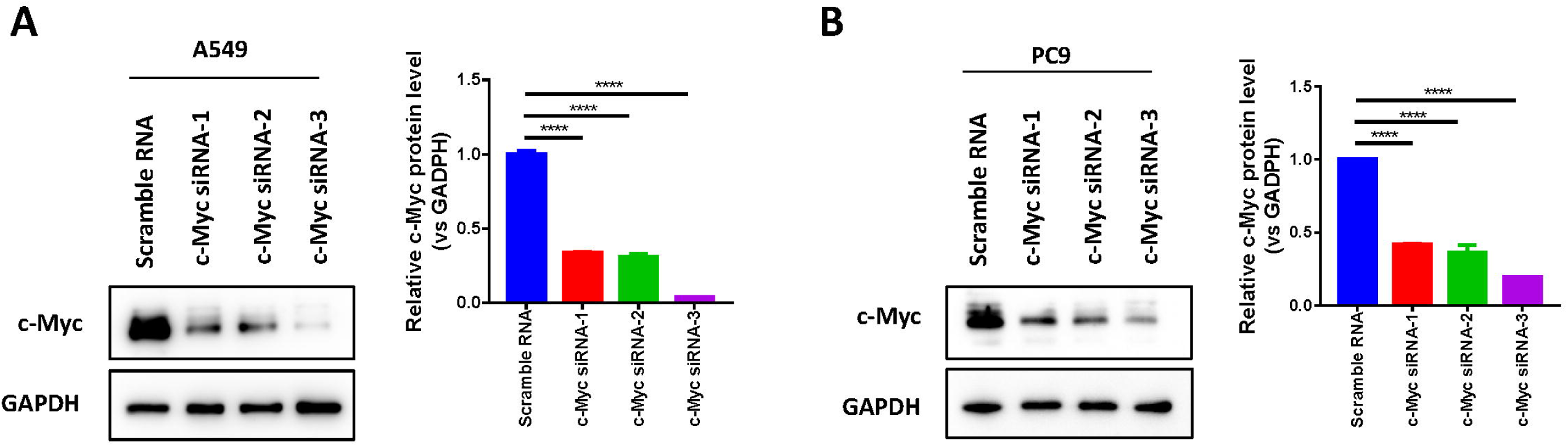

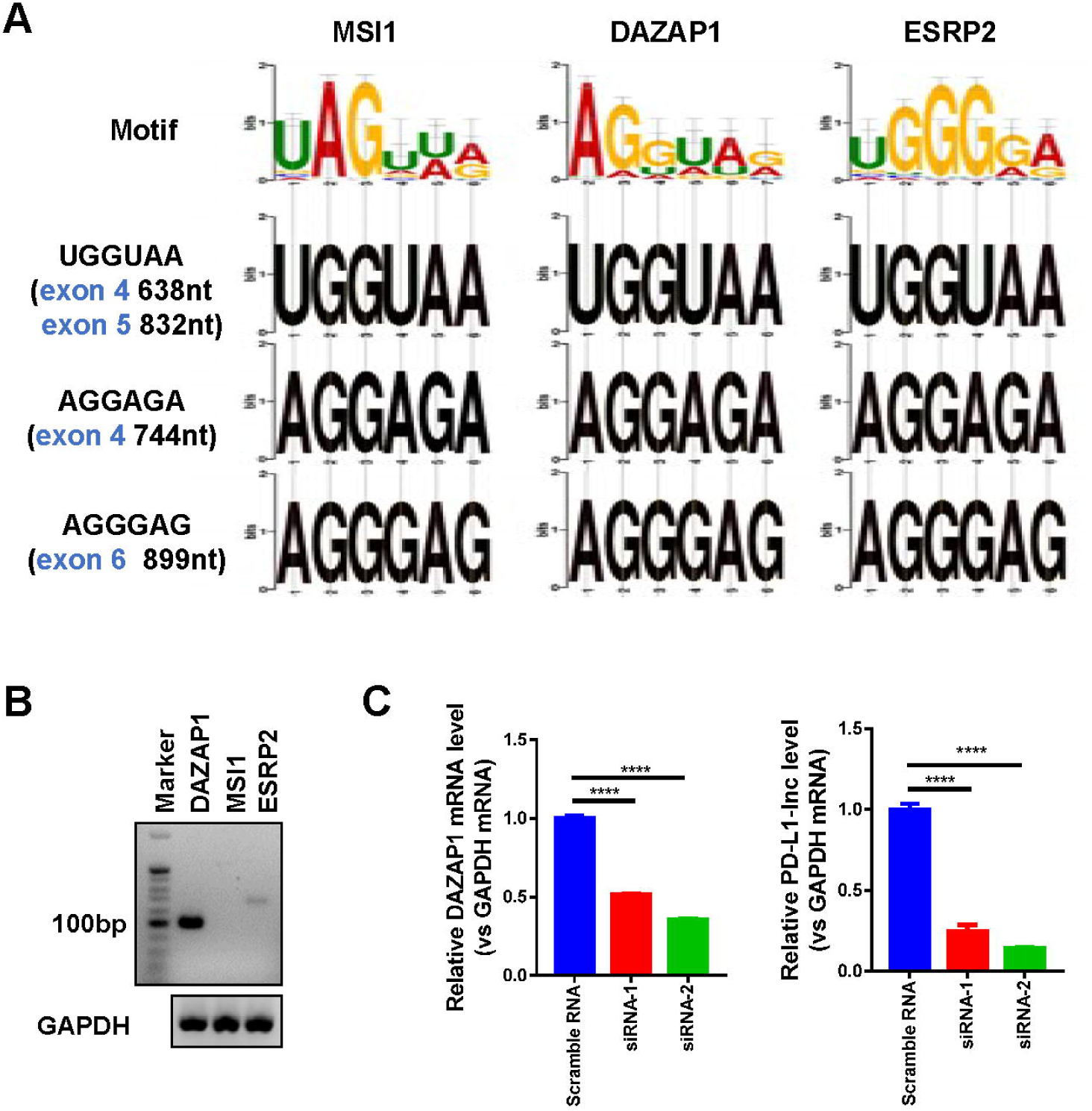

## Notes

### Competing Interest Statement

The authors have declared no competing interest.

