## Supplementary material for "PD-L1 lncRNA splice promotes lung adenocarcinoma progression via enhancing c-Myc activity": manuscripts

**File list**

1. Table S1
2. Table S2
3. Table S3
4. Figure S1
5. Figure S2
6. Figure S3
7. Figure S4
8. Figure S5
9. Figure S6
10. Figure S7
11. Figure S8
12. Figure S9
13. Figure S10
14. Figure S11

**Table S1. Clinic characteristics of lung adenocarcinoma patients.**

| | PD-L1 Negative | PD-L1 Positive | P value |
|---|---|---|---|
| **Parameter** | | | |
| **All patients** | N=212 (77.1%) | N=63 (22.9%) | |
| **Age (years)** | | | **0.827** |
| <60 | 84 (77.8%) | 24 (22.2%) | |
| ≧60 | 128 (76.6%) | 39 (23.4%) | |
| **Gender** | | | |
| Female | 136 (85%) | 24 (15%) | **0.000** |
| Male | 76 (66%) | 39 (34%) | |
| **Pathologic stage** | | | |
| I | 164 (82.4%) | 35 (17.6%) | **0.001** |
| II-III | 48 (70.6%) | 28 (29.4%) | |
| **Pathologic T stage** | | | |
| T1 | 142 (82.6%) | 30 (17.4%) | **0.005** |
| T2-T4 | 70 (68.0%) | 33 (32.0%) | |
| **Pathologic N stage** | | | |
| N0 | 171 (82.2%) | 37 (17.8%) | **0.000** |
| N1-N3 | 41 (62.0%) | 26 (38.0%) | |

**Table S2. List of primers for qRT-PCR.**

| Gene name | Primers | |
|---|---|---|
| | Forward | Reverse |
| **GAPDH** | TGAACGGGAAGCTCACTGG | TCCACCACCCTGTTGCTGTA |
| **PD-L1 mRNA primer 1** | AGGCCGAAGTCATCTGGAC | CTGGGATGACCAATTCAGCT |
| **PD-L1 mRNA/lnc primer 2** | TCACGGTTCCCAAGGACCTA | CCCCGATGAACCCCTAAACC |
| **PD-L1 mRNA primer 3** | CCAGTCACCTCTGAACATG | TCAGTGTGCTGGTCACATTG |
| **PD-L1-lnc primer 4** | CTGAGTGGAGATTAGATCCTG | CATCATTCTCCCAAGTGAGTC |
| **SAA2** | GCTTCTTTTCGTTCCTTGGCG | GCCGATGTAATTGGCTTCTCTCA |
| **OAS2** | ACGTGACATCCTCGATAAAACTG | GAACCCATCAAGGGACTTCTG |
| **IFI44L** | ACAGAGCCAAATGATTCCCTATG | TCGATAAACGACACACCAGTTG |
| **SAA4** | GGCAGAGCCTATTGGGACATA | GCTGATGAGTTTAGCAGCCC |
| **IFI44** | ATGGCAGTGACAACTCGTTTG | TCCTGGTAACTCTCTTCTGCATA |
| **OASL** | CTGATGCAGGAACTGTATAGCAC | CACAGCGTCTAGCACCTCTT |
| **IFITM1** | CCAAGGTCCACCGTGATTAAC | ACCAGTTCAAGAAGAGGGTGTT |
| **MX2** | CAGAGGCAGCAGACGATCAAC | TTGGTCAGGATACCGATGGTC |
| **MX1** | GTTTCCGAAGTGGACATCGCA | CTGCACAGGTTGTTCTCAGC |
| **COL8A1** | GCTGCCACCTCAAATTCCTC | CTTTCTTGGGTACGGCTTCCT |
| **IFIT1** | AGAAGCAGGCAATCACAGAAAA | CTGAAACCGACCATAGTGGAAAT |
| **IFIT3** | AAAAGCCCAACAACCCAGAAT | CGTATTGGTTATCAGGACTCAGC |
| **SOD2** | GCTCCGGTTTTGGGGTATCTG | GCGTTGATGTGAGGTTCCAG |
| **IFITM3** | ACTGTCCAAACCTTCTTCTCTCC | TCGCCAACCATCTTCCTGTC |
| **IFI6** | GGTCTGCGATCCTGAATGGG | TCACTATCGAGATACTTGTGGGT |
| **HLA-A** | GACGCCCCCAAAACGCATA | TGGGCAAACCCTCATGCTG |
| **GFP mRNA** | AAGGACGACGGCAACTACAA | CGATGTTGTGGCGGATCTTG |
| **MSI1** | GGGACTCAGTTGGCAGACTAC | CTGGTCCATGAAAGTGACGAA |
| **ESRP2** | TTGCAGCAAGGCTGATGTG | GTTGAGGCAGAGTGCTACACC |
| **DAZAP1** | AGAAGTTCGGAGTGGTCACG | ACTGATTGTTCGTCCTCGAAAG |

**Table S3. List of target sequences of various siRNAs.**

| Gene name | Target sequence 5'-3' |
|---|---|
| **PD-L1-lnc shRNA** | GACTCACTTGGGAGAATGATGGA |
| **c-Myc-si-1** | GAGGAGACATGGTGAACCA |
| **c-Myc-si-2** | GGGTCAAGTTGGACAGTGT |
| **c-Myc-si-3** | CGACGAGACCTTCATCAAA |
| **DAZAP-si-1** | CGGAGGTAGTCATGATCTA |
| **DAZAP-si-2** | CAACCAGTCTCGAGGCTTT |

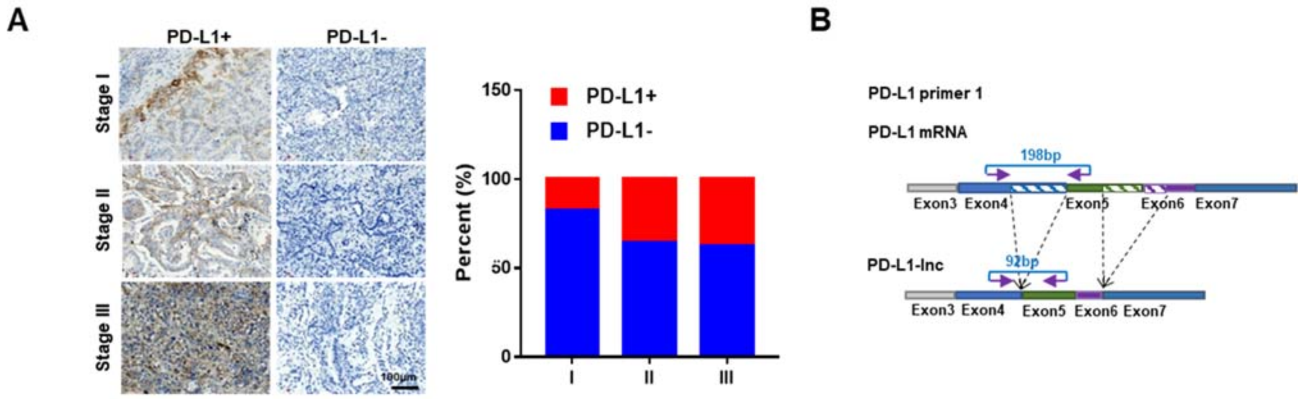


**Figure S1. The PD-L1 protein level in human lung cancer tissue. (A)** Immunohistochemical staining for PD-L1 in human lung cancer tissue sections. Left: representative image; Right: quantitative analysis. Tumors were classified into three stages (Stage I, II and II) according to the 7th edition of the American Joint Committee on Cancer Staging Manual 17. **(B)** The schematic of primer for amplification of PD-L1 mRNA.

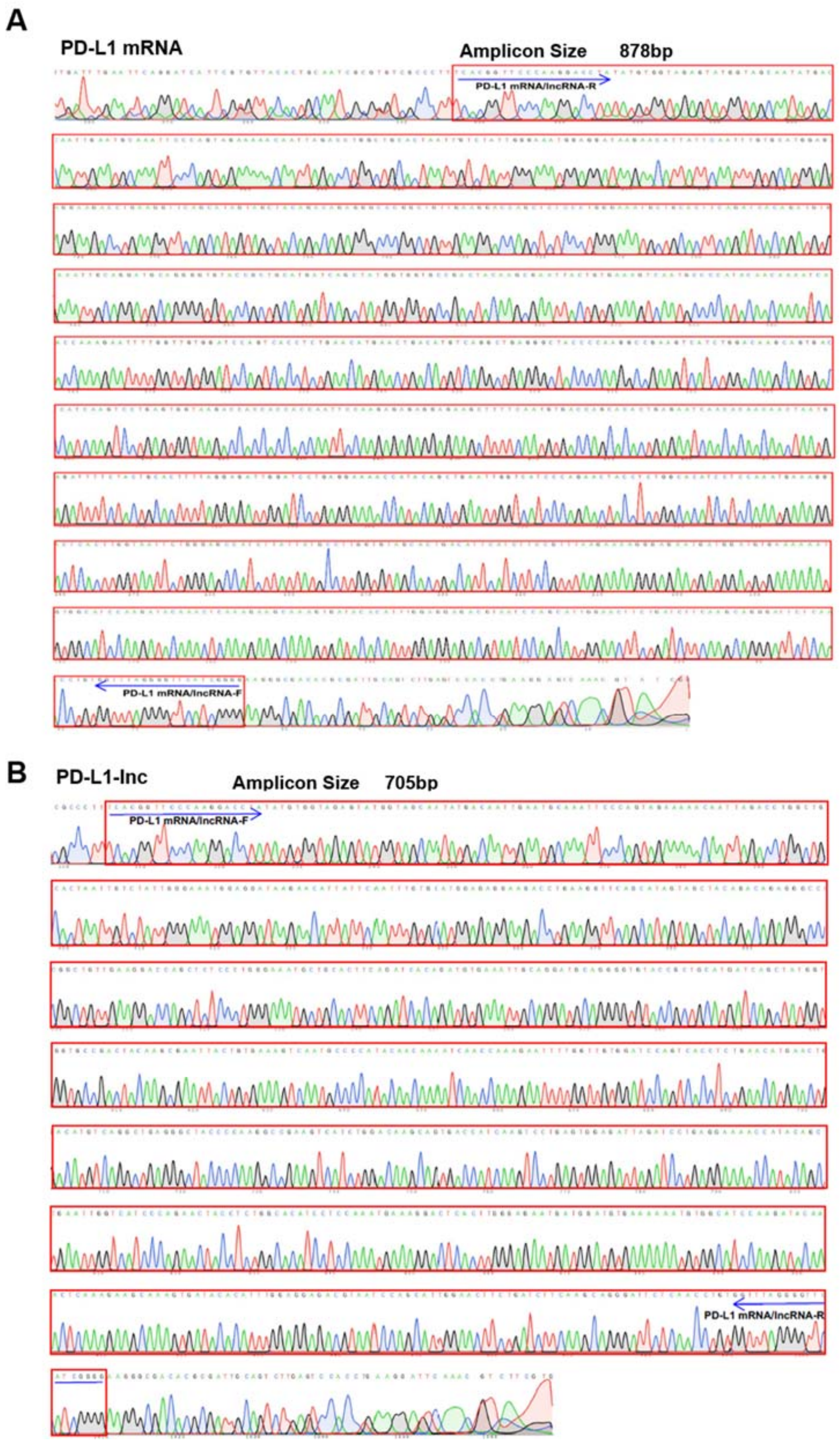


**Figure S2.** The Sanger sequencing result of the (A) PD-L1 mRNA (878*bp* band) and (B) PD-L1 lncRNA (705*bp* band) obtained by RT-PCR.

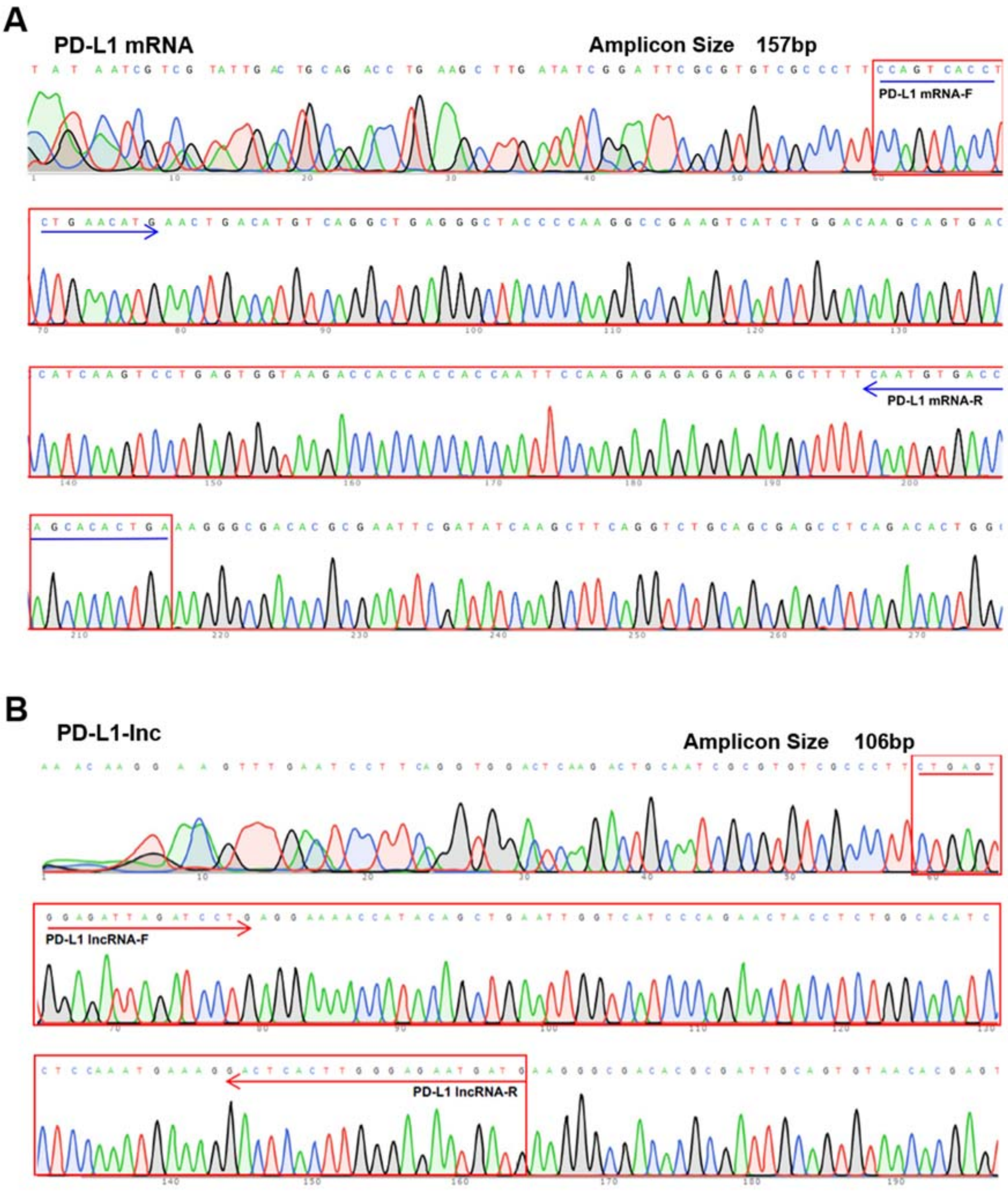


**Figure S3.** The Sanger sequencing result of the PD-L1 mRNA obtained by qRT-PCR with specific primer for amplification of (A) PD-L1 mRNA (157*bp* band) and (B) PD-L1 lncRNA (106*bp* band) obtained by RT-PCR.

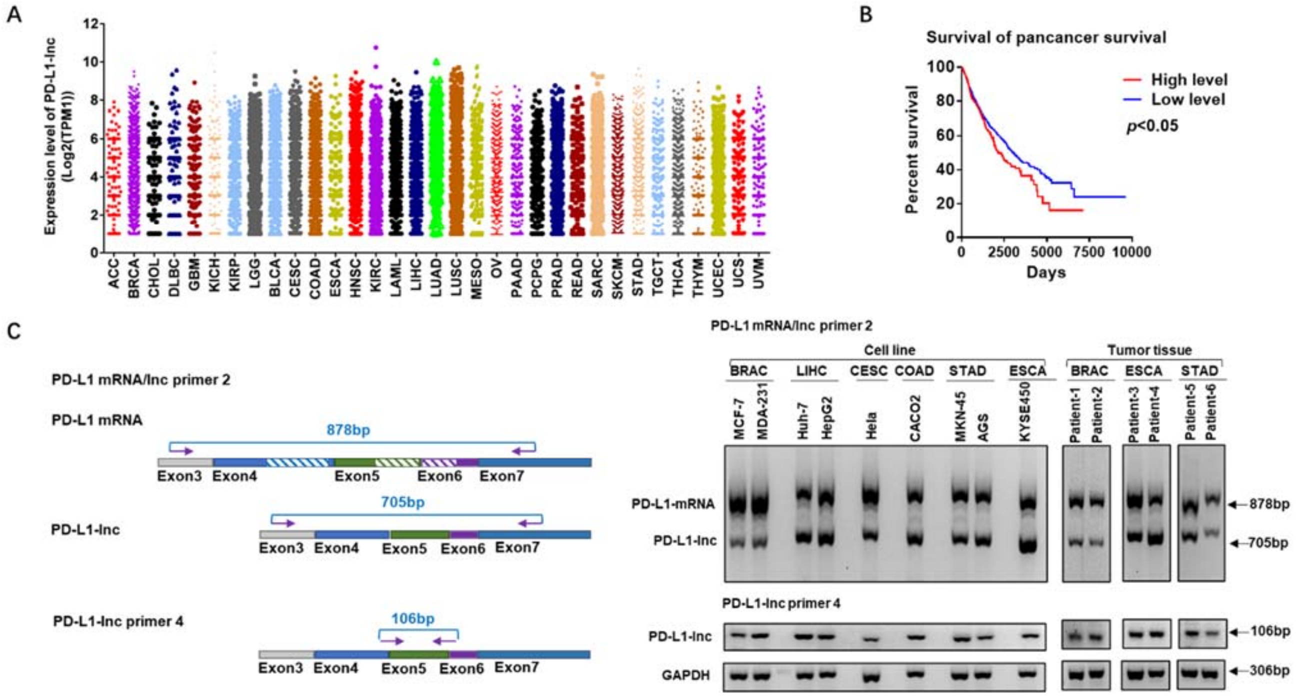


**Figure S4. Expression of PD-L1-lnc in various cancers. (A)** Pan-cancer analysis of PD-L1-lnc expression in TCGA database. **(B)** PD-L1-lnc as predictors of cancer survival in TCGA database. **(C)** Left, the schematic of primer for amplification of PD-L1 mRNA and PD-L1-lnc; Right, the agarose gel of the PD-L1 mRNA and PD-L1-lnc in BRAC, LIHC, CESC, COAD, STAD and ESCA cancer cell lines and BRAC, ESCA and STAD cancer tissues by RT-PCR with specific probes. **ACC**: Adrenocortical carcinoma; **BRCA**: Breast invasive carcinoma; **CHOL**: Cholangiocarcinoma; **DLBC**: Lymphoid neoplasm diffuse large B-cell lymphoma; **GBM**: Glioblastoma multiforme; **KICH:** Kidney chromophobe; **KIRP**: Kidney renal papillary cell carcinoma; **LGG**: Brain lower grade glioma; **BLCA**: Bladder urothelial carcinoma; **CESC**: Cervical squamous cell carcinoma and endocervical adenocarcinoma; **COAD**: Colon adenocarcinoma; **ESCA**: Esophageal carcinoma; **HNSC**: Head and neck squamous cell carcinoma; **KIRC**: Kidney renal clear cell carcinoma; **LAML**: Acute myeloid leukemia; **LIHC:** Liver hepatocellular carcinoma; **LUAD**: Lung adenocarcinoma; **LUSC**: Lung squamous cell carcinoma; **MESO**: Mesothelioma; **OV**: Ovarian serous cystadenocarcinoma; **PAAD**: Pancreatic adenocarcinoma; **PCPG**: Pheochromocytoma and paraganglioma; **PRAD**: Prostate adenocarcinoma; **READ**: Rectum adenocarcinoma; **SARC**: Sarcoma; **SKCM**: Skin cutaneous melanoma; **STAD**: Stomach adenocarcinoma; **TGCT**: Testicular germ gell tumors; **THCA**: Thyroid carcinoma; **THYM:** Thymoma; **UCEC:** Uterine corpus endometrial earcinoma; **UCS**: Uterine carcinosarcoma; **UVM**: Uveal Melanoma

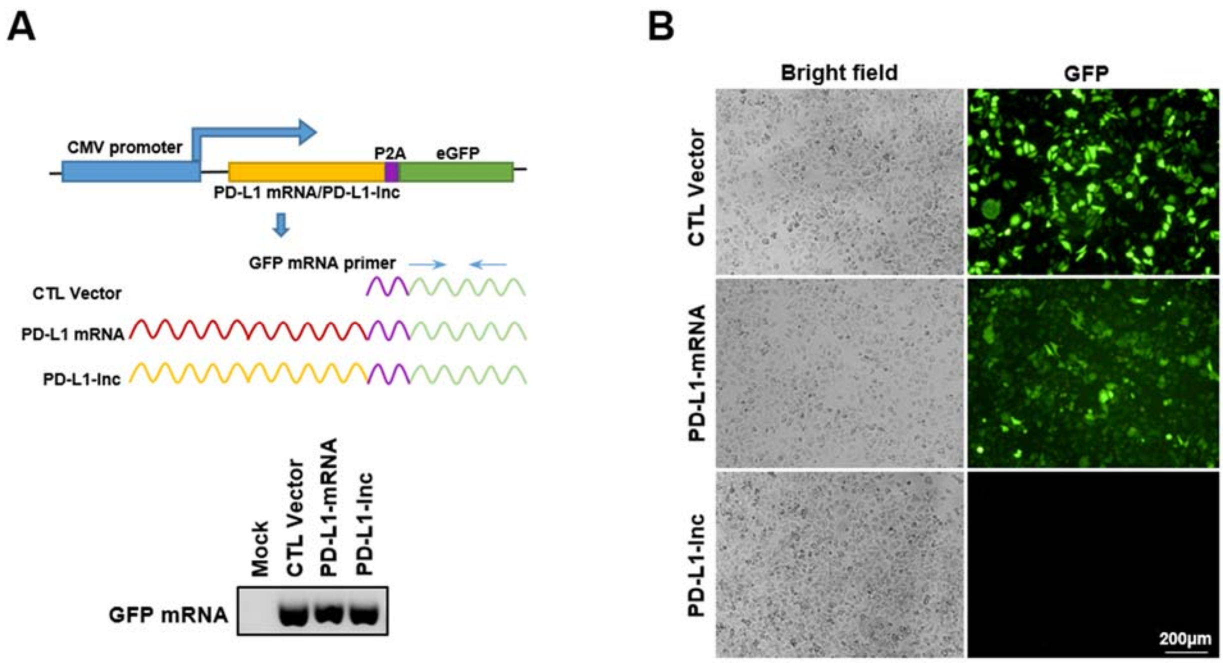


**Figure S5. PD-L1-lnc is a long non-coding RNA fragment. (A)** Linking PD-L1-lnc or PD-L1-mRNA with GFP mRNA to generate the 'recombinant' GFP expression vector. **(B)** GFP expression detected by fluorescence microscopy in A549 cells than were transfected with GFP expression or 'recombinant' GFP expression vectors. PD-L1-lnc overexpression vector: PD-L1-lnc; PD-L1 overexpression vector: PD-L1-mRNA; control vector: CTL Vector.

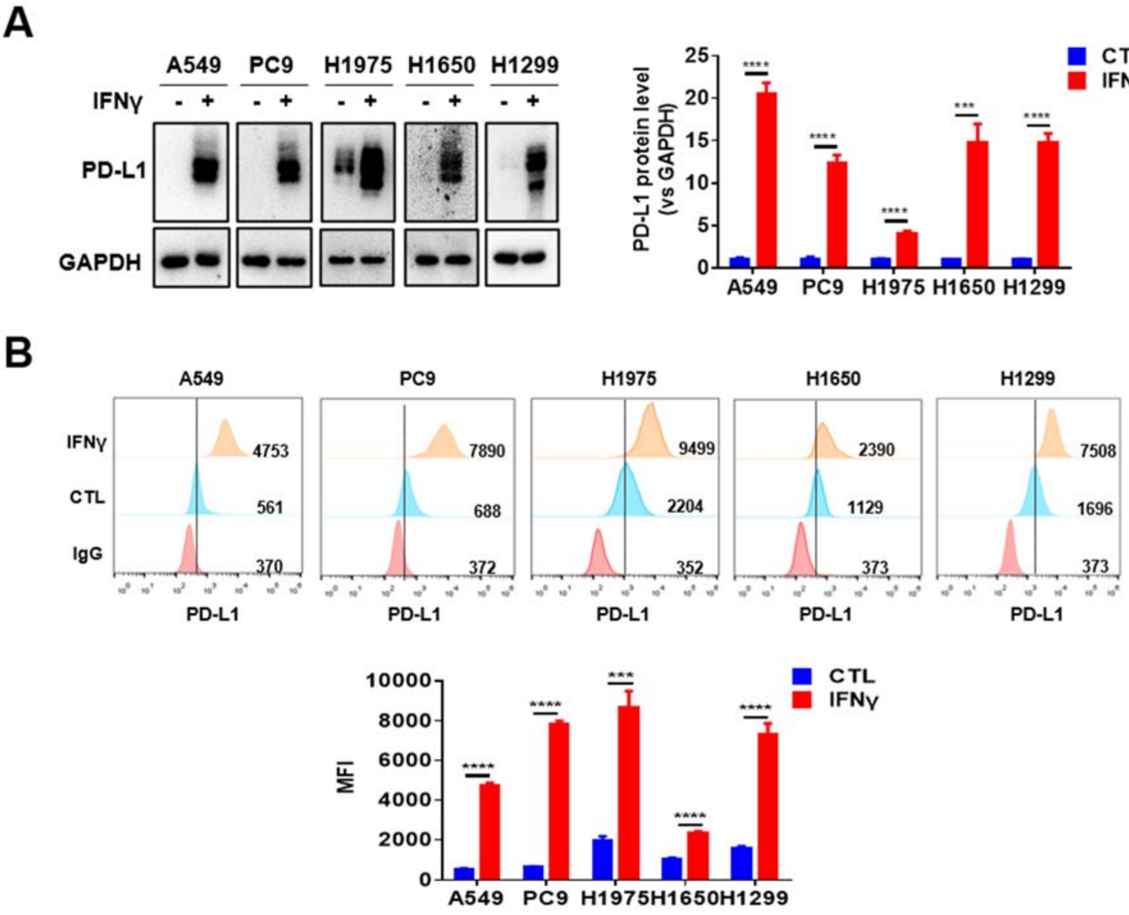


**Figure S6.** Induction of PD-L1 by IFNγ in lung adenocarcinoma cells. **(A)**The expression level of PD-L1 protein in lung adenocarcinoma cells with or without IFNγ stimulation determined by Western blotting. **(B)** The expression level of PD-L1 protein in lung adenocarcinoma cells with or without IFNγ stimulation determined by flow cytometry. In a-b, left: representative image; right: quantitative analysis. ***$P < 0.001$; ****$P < 0.0001$.

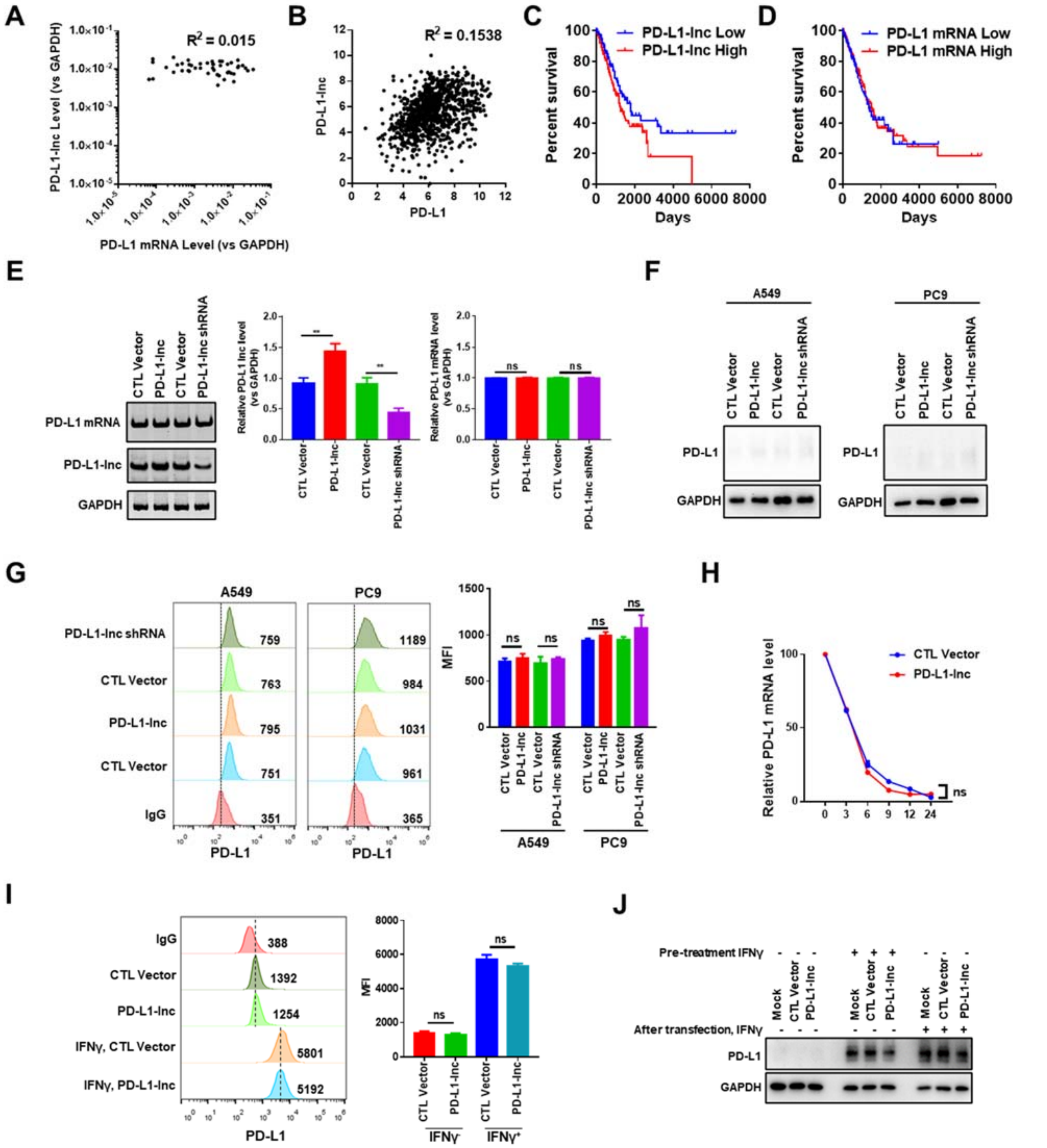


**Figure S7. PD-L1-lnc has no effect on PD-L1 mRNA and protein expression. (A)** Relative expression of the PD-L1-lnc and PD-L1 mRNA in PD-L1 positive and negative lung adenocarcinoma tissues. **(B)** Relative expression of the PD-L1-lnc and PD-L1 mRNA in lung adenocarcinoma data set of TCGA. **(C-D)** Kaplan–Meier survival analysis of lung adenocarcinoma patients stratified by PD-L1-lnc (C) and PD-L1 mRNA (D) expression in TCGA database. **(E)**The expression of PD-L1 lncRNA and mRNA in A549 cells transfected with PD-L1-lnc overexpression vector or PD-L1-lnc shRNA vector. **(F-G)** wester blotting (F) and Flow cytometry

(G) analysis of PD-L1 protein level in A549 and PC9 cells transfected with PD-L1-lnc overexpression vector or PD-L1-lnc shRNA vector. **(H)** The half-life of PD-L1 mRNA in A549 cells transfected with PD-L1-lnc overexpression vector or control vector. **(I-J)** The expression of PD-L1 mRNA (I) and protein (J) levels in A549 cells transfected with PD-L1-lnc overexpression vector or control vector in the presence or absence of IFNγ. PD-L1-lnc overexpression vector: PD-L1-lnc; control vector: CTL Vector. ** $P < 0.01$.

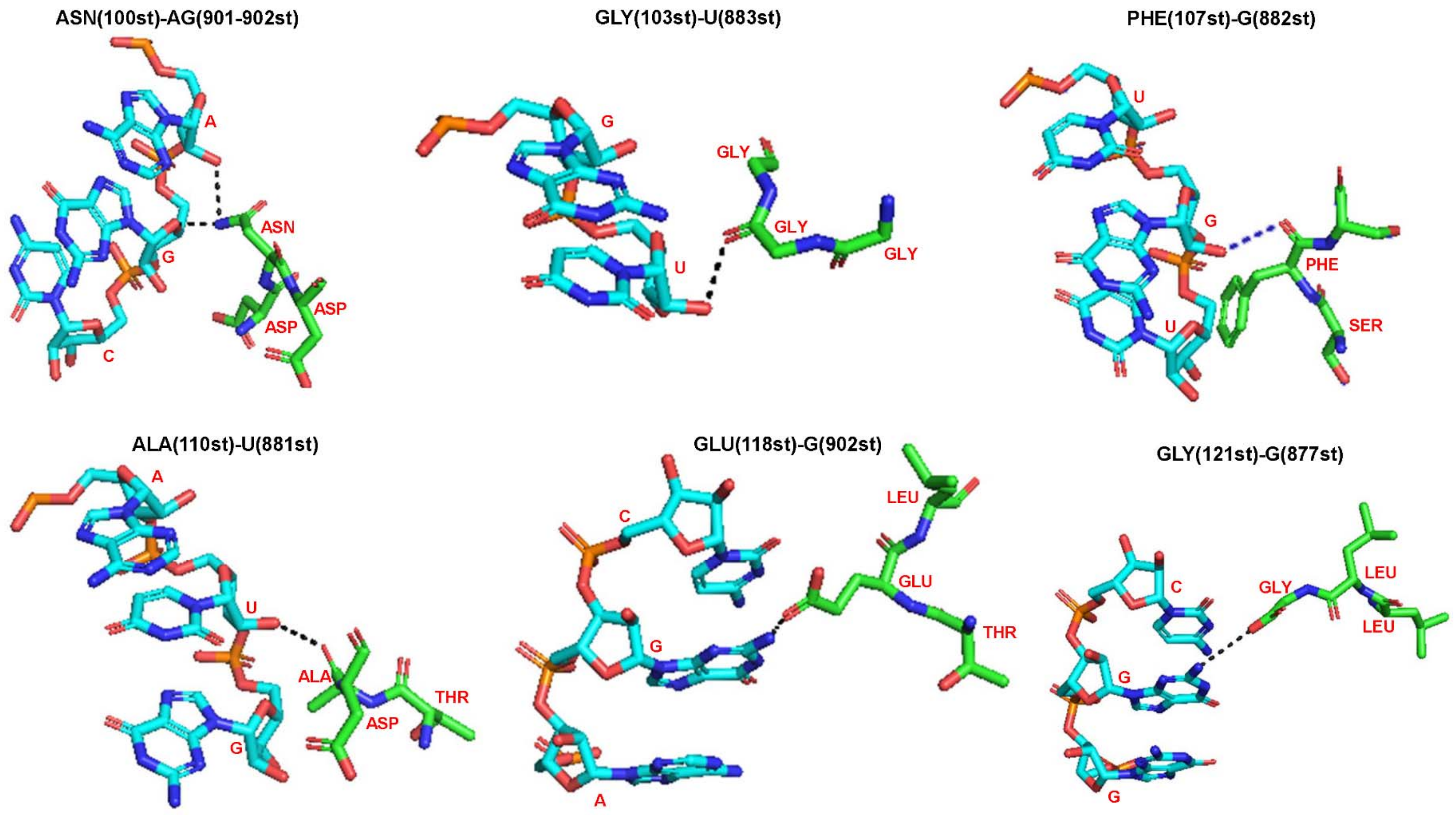


**Figure S8. Graphical representation images of the binding interface of the docking models of PD-L1-lnc with c-Myc by NPDock.** Black dashes represent hydrogen bond

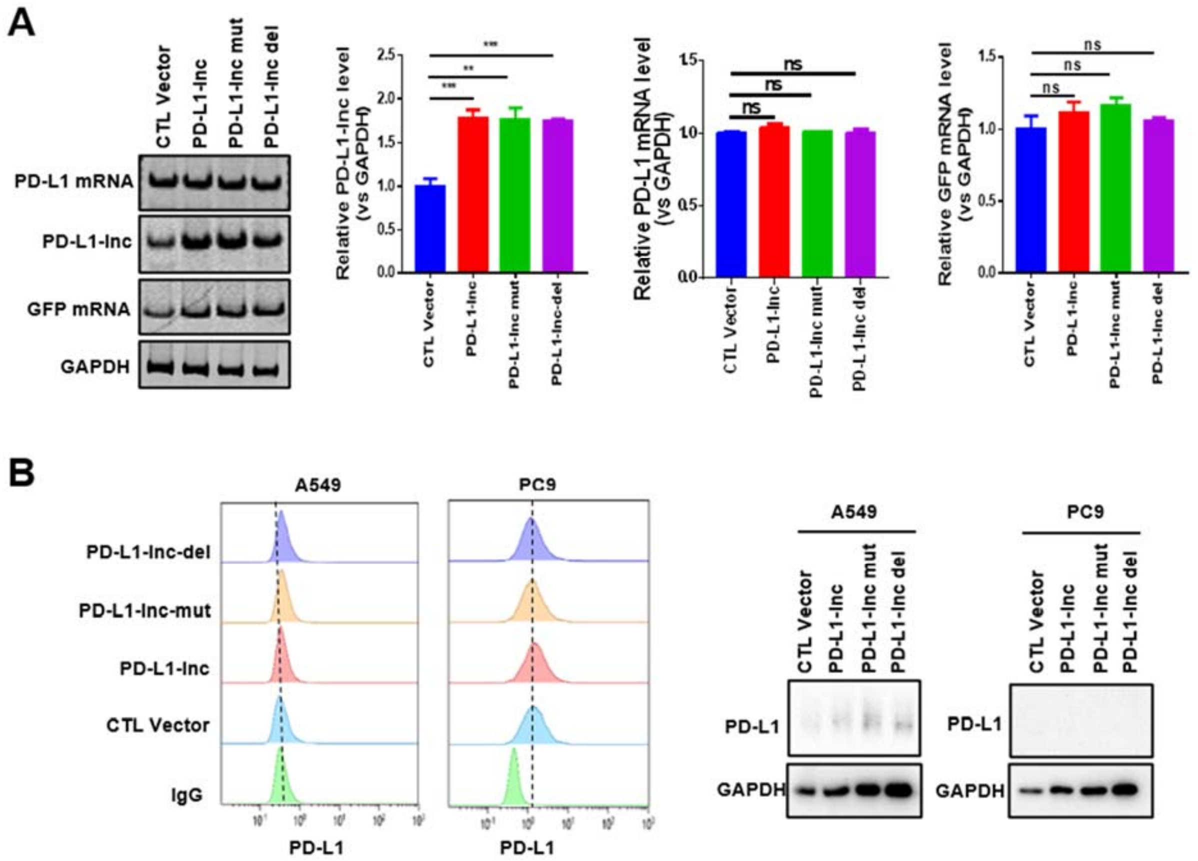


**Figure S9. The expression of PD-L1 lncRNA and mRNA in A549 cells transfected with the mutant vector and wild type vector. (A)** Agarose gel analysis of the expression of PD-L1 lncRNA and mRNA in A549 cells transfected with the mutant vectors and wild type vector. **(B)** Flow cytometry (left) and western blotting (right) analysis of the expression of PD-L1 protein in A549 cells transfected with the mutant or wild type PD-L1 vector. PD-L1-lnc overexpression vector: PD-L1-lnc; control vector: CTL Vector. ***P* < 0.01; ****P* < 0.001.

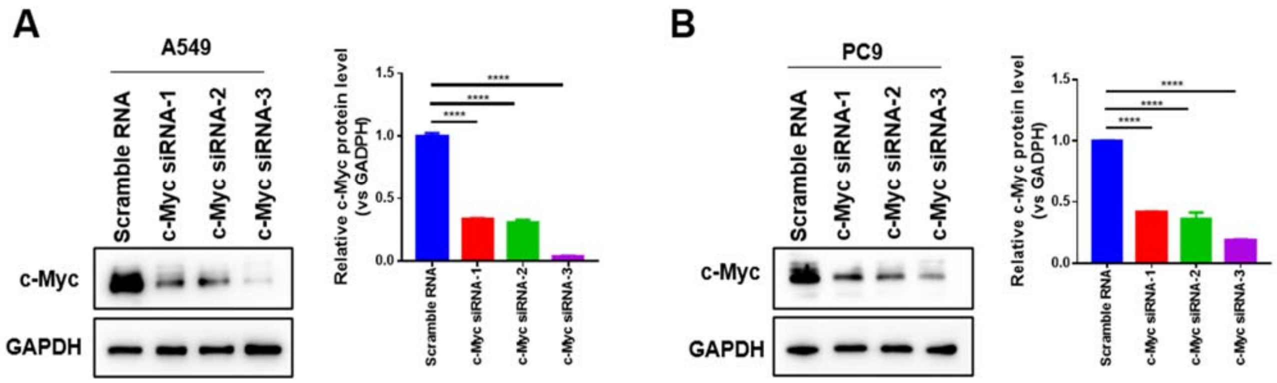


**Figure S10.** The efficiency of c-Myc siRNA in A549 (A) and PC9 (B) by Western blotting. ****$P < 0.0001$.

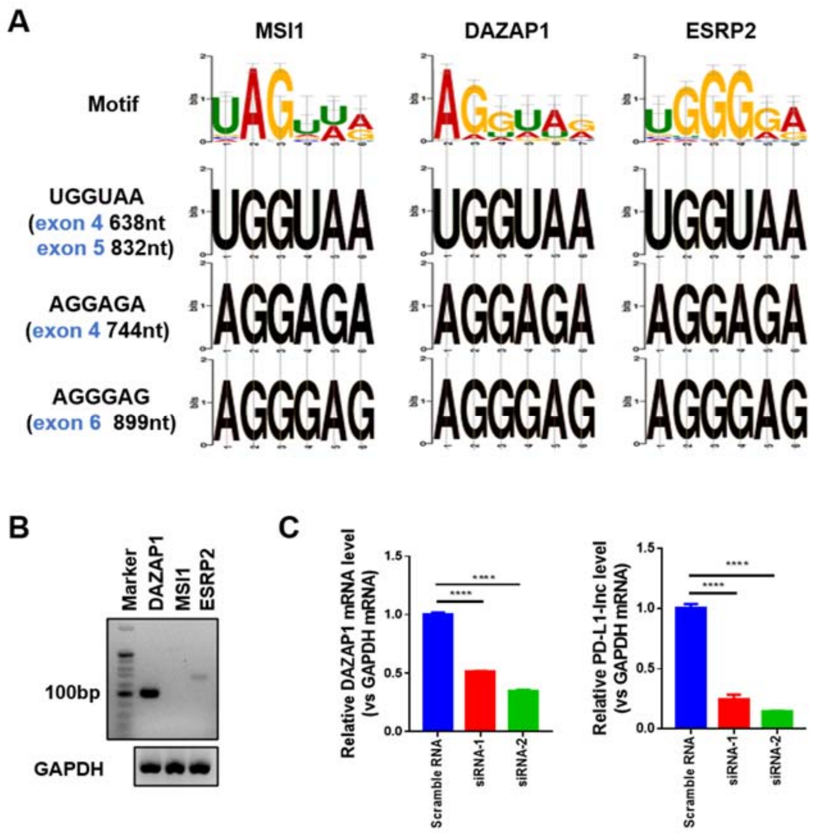


**Figure S11. The effect of DAZAP1 on the generation of PD-L1-lnc. (A)** The motif of MSI1, DAZAP1 and ESRP2 and alternative splicing site of PD-L1. **(B)** The efficiency of DAZAP1 siRNA on DAZAP1 reduction. **(C)** The expression level of DAZAP1 mRNA (left) and PD-L1-lnc (right) in A549 cells transfected with siRNAs to DAZAP1. *****P* < 0.0001.
